# The condensation of HP1-α/Swi6 imparts nuclear stiffness

**DOI:** 10.1101/2020.07.02.184127

**Authors:** Jessica F. Williams, Ivan V. Surovtsev, Sarah M. Schreiner, Ziyuan Chen, Gulzhan Raiymbek, Hang Nguyen, Yan Hu, Julie S. Biteen, Simon G.J. Mochrie, Kaushik Ragunathan, Megan C. King

## Abstract

The condensation of proteins and nucleic acids underlies the formation of membraneless organelles, which have emerged as major drivers of cellular organization. It remains largely unexplored, however, whether these condensates can impart mechanical function(s) to the cell. The heterochromatin protein HP1-α (Swi6 in *S. pombe*) crosslinks histone H3K9 methylated nucleosomes and has been proposed to undergo condensation to drive the liquid-like clustering of heterochromatin domains. Here we leverage the genetically tractable *S. pombe* model and a separation-of-function Swi6 allele to elucidate a mechanical function imparted by its condensation. Using a combination of single-molecule imaging, force spectroscopy on individual nuclei, and high-resolution live-cell imaging, we show that Swi6 is critical for nuclear resistance to external force. Strikingly, it is this condensed yet dynamic pool of Swi6, rather than the chromatin-bound molecules, that is essential to imparting mechanical stiffness. Our findings suggest that Swi6 condensates embedded in the chromatin meshwork establish the emergent mechanical behavior of the nucleus as a whole, revealing that biomolecular condensation can influence organelle and cell mechanics.

## INTRODUCTION

The cell nucleus provides a mechanically resilient compartment that encapsulates and protects the genome^1,2^. The nuclear compartment is defined by the nuclear envelope, an integrated network of the inner and outer nuclear membranes, nuclear pore complexes, integral inner nuclear membrane proteins, peripheral chromatin, and, in higher eukaryotes, a polymer mesh of intermediate filaments (lamins)^3–5^. Maintaining the integrity of the nuclear envelope is critical for cell survival, and mechanical defects that lead to nuclear blebbing and rupture are tied to human disease^1,2^. Nuclear ruptures can occur during cellular migration, when cells move through tissue and experience significant compressive force^6,7^, but even intrinsic cytoskeletal forces in adherent or contractile cells are strong enough to induce nuclear rupture, indicating that the nucleus is under constant strain from its own cellular architecture^8,9^.

In the effort to understand how the nucleus withstands these mechanical forces, much work has focused on the contribution of the nuclear lamina^10,11^. However, we previously demonstrated that the tethering of chromatin to the nuclear envelope is critical for maintaining nuclear stiffness in fission yeast, which lack nuclear lamins^12^. Other studies in mammalian cells have further demonstrated the unique mechanical role of chromatin in its contribution to the nuclear force response to small deformations (1-3 μm), while A-type lamins are critical for the force response to large deformations (> 3 μm)^13^. Additionally, when chromatin compaction is globally altered through histone deacetylase or demethylase inhibition, the relative proportion of euchromatin to heterochromatin (HC) affects the overall stiffness of the nucleus^14^. Thus, both the lamina and the underlying chromatin are important for maintaining mechanical stability, yet each play separate roles related to the physical—and perhaps temporal—scale of the external strain. Indeed, increasing the amount HC is sufficient to rescue the nuclear irregularity and nuclear blebbing phenotype seen in HeLa cells expressing the mutant form of lamin A (progerin) of the premature aging disease Hutchinson-Guilford Progeria Syndrome^14^. Together, these studies suggest that in addition to its function as a transcriptional insulator^15^, HC acts as a dynamic mechanical scaffold for the nucleus^16^; however, the biophysical mechanisms by which HC can influence nuclear mechanics remain largely undefined.

An evolutionarily conserved feature of nuclear organization is the association of HC with the nuclear periphery. This localization has classically been thought to reflect the ability of HC to promote gene silencing. Several studies, however, suggest that the causality of the periphery-silencing relationship may be more complex, implying that there may be additional function(s) for peripherally-located HC. Indeed, in *C. elegans*, HC positioning at the nuclear periphery does not dictate gene silencing^17^. Moreover, leveraging the model of rod cells with “inverted” nuclei, re-localization of HC from the periphery to the center of the nucleus was found to neither alter gene expression nor chromatin organization at the 100s of kilobase to many megabase scale^18^. Our previous work demonstrated that chromatin tethering to the nuclear periphery is important for resisting nuclear deformations in response to microtubule forces in fission yeast^12,19^, which is echoed by studies in mammalian cells^14,20^. Here we further probe the hypothesis that a primary function of peripherally-located HC is to mechanically support the nuclear envelope.

The potential mechanism for this mechanical support stems from recent evidence suggesting that a key protein responsible for HC compaction and gene silencing— heterochromatin protein (HP1-α)—is important for maintaining normal nuclear morphology and stiffness without affecting HC peripheral localization^21^ and forms phase-separated, HP1α-rich HC domains^22,23^. These (and other) biomolecular condensates often exhibit liquid-like properties, including surface tension and viscosity^24–29^. One model proposes that the surface tension of HC-associated condensates, enhanced by strong electrostatic interactions as observed for HP1-α^30^, would resist deformation. Alternatively, or in addition, a distinct model of “elastic ripening” is being developed^31^, in which the formation of condensates embedded in a polymer network gives rise to a composite system with emergent or synergistic mechanical properties. Lastly, while it has been proposed that biomolecular condensates can do mechanical work^32^, for example, to drive local membrane bending^33–35^ or, in an artificial system, to bring two chromatin loci together^36^, whether biomolecular condensation can drive emergent mechanical behaviors at the scale much larger than an individual condensate, i.e., at the level of organelles and cells, has yet to be explored.

To biomechanically define the features of HP1-α/Swi6 HC domains, we employed both *in vivo* fluorescence microscopy and *in vitro* force spectroscopy techniques. We demonstrate that HC domains respond to mechanical force with viscoelastic properties, and that proper coalescence of the domains—specifically telomeres—depends on HP1-α/Swi6. Using single-molecule imaging we describe a separation-of-function allele of HP1-α/Swi6—*swi6-sm1*—which displays disruption of the biomolecular condensation of HC domains despite normal H3K9me2/3 binding and crosslinking *in vivo*. This defect in Swi6 condensation leads to depletion of mobile Swi6 at HC domains and concomitantly leads to substantially softer nuclei both *in vitro* and *in vivo*. Taken together, our findings argue that it is the condensation of HP1-α/Swi6 at HC domains, rather than the binding and/or crosslinking of H3K9me2/3 nucleosomes, that contributes to the overall stiffness of the nucleus, showcasing a role for biomolecular condensates within a biological meshwork (the chromatin) in driving the emergent mechanical stiffness of a membrane-bound compartment.

## RESULTS

### Heterochromatin domains exhibit liquid-like properties *in vivo* that respond to cytoskeletal forces

The HC protein HP1-α/Swi6 is a key component of centromeric and telomeric HC, responsible for its condensation and transcriptional repression^37–39^ and has been shown to oligomerize and form phase-separated domains that facilitate HC formation^22,23,40^. With this in mind, we sought to further characterize the liquid-like properties of Swi6 at two regions at which it is highly enriched: the centromeres and the telomeres^37,41,42^.

When Swi6 is visualized as a fluorescent protein fusion expressed from its endogenous locus (Swi6-GFP), it resides in several nuclear foci/domains (Figure 1A, top). The largest domain is comprised of the centromeres of the three chromosomes, which are coupled to the spindle-pole body (SPB) interface (Figure 1A, bottom). The other, smaller heterochromatic foci contain the six telomeres, the mating type locus, and other silenced regions of the genome and are typically associated with the nuclear envelope far from the SPB. In *S. pombe*, microtubules continuously translocate the SPB, which transmits the MT-dependent force on to the centromeric HC. As the centromeric HC is driven across the nucleus by MT forces, we routinely observe a morphologic transition from a circular domain (as visualized from a two-dimensional image plane; Figure 1B) when “at rest” (at t = 0) to an elliptical domain when under maximum MT shearing force (at t = 30-80s), stretching along the SPB oscillation axis (Figure 1B; see also insets of reconstructed contours in green). At various time points during this translational motion (t > 60s), the centromere displays an apparent fission into smaller, individual domains that later merge (t > 100s). These observations suggest that mechanical forces from the cytoskeleton are directly transduced to HC, leading to changes in domain morphology consistent with liquid-like reorganization^26,43^. We therefore considered the possibility that HC domains can act as mechanical cushions, absorbing MT-driven forces exerted upon the nucleus through their resistance to deformation and/or their effects on the emergent properties of the chromatin itself.

**Figure 1.**
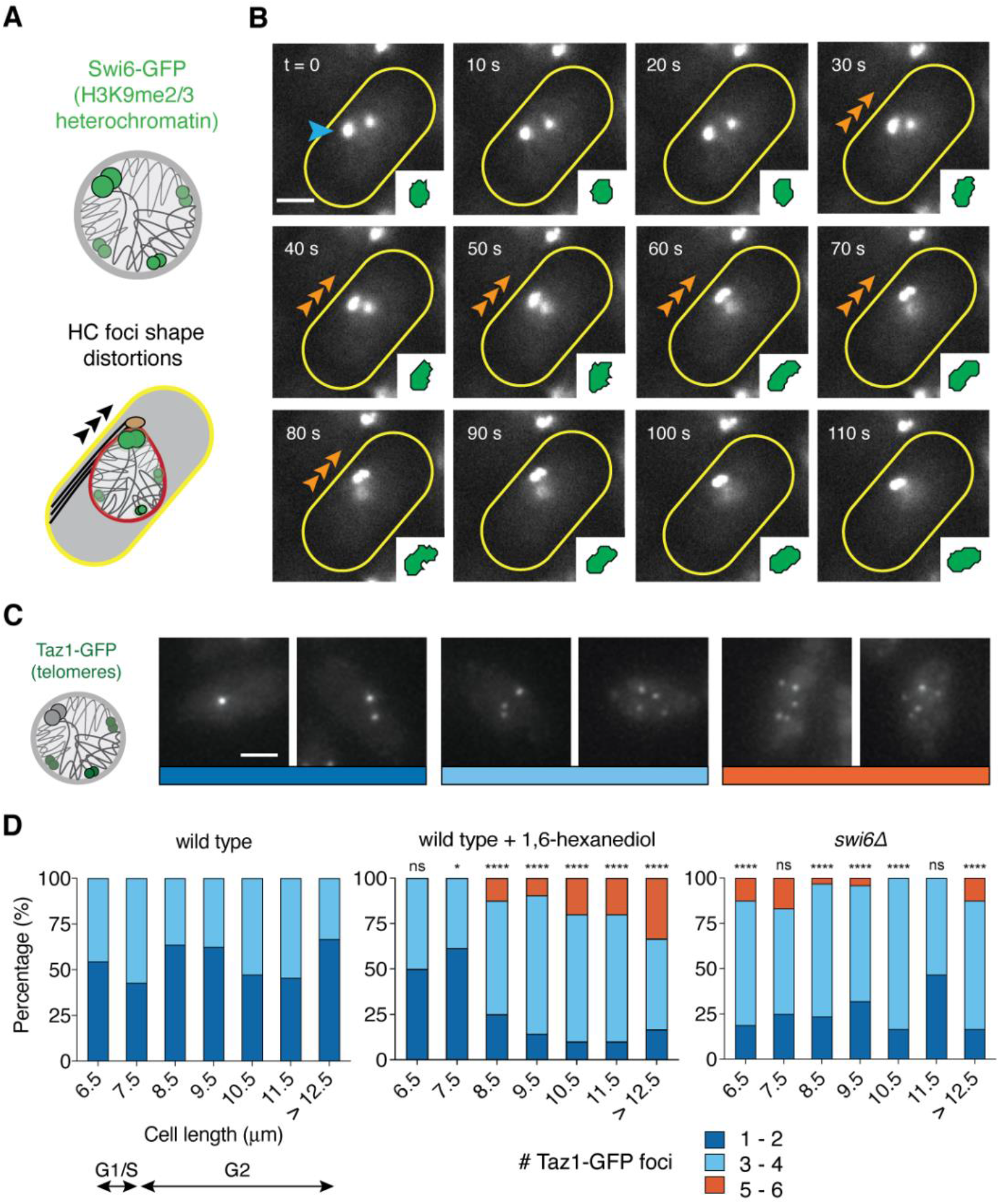
Heterochromatin domains exhibit liquid-like properties *in vivo* that are tied to normal telomere clustering. (A) Schematic representation of the distribution of heterochromatin (HC) domains in *S. pombe* as visualized by expression of Swi6-GFP (light green) from its endogenous locus. HC domains are subject to shearing forces driven by microtubule dynamics. (B) HC domains are deformed in response to MT-driven nuclear oscillations. Time-lapse montage of cells expressing Swi6-GFP shows a large HC focus (blue arrowhead) transitioning from a round to an oblong shape that aligns with the long axis of the cell as it is translated in response to MT dynamics (orange arrowheads, t = 30 to t = 80 s); image insets display the contour of thresholded Swi6-GFP foci. At t = 60 s, the focus begins to separate into two foci, and after MT dynamics cease (t = 100 s), these foci begin to coalesce. Scale bar, 2 μm. (C,D) Under normal conditions, the six fission yeast telomeres typically cluster into ∼1-3 foci during interphase that de-cluster in response to either 1,6-hexanediol treatment or deletion of Swi6. (C) Fluorescence maximum intensity projections of representative *S. pombe* cells expressing Taz1-GFP with 1-6 telomere clusters (see also Figure S2). Scale bar, 2 μm. (D) Percentage of cells containing the indicated number of Taz1-GFP foci; foci per cell were counted and grouped by cell length as a proxy for cell cycle progression. In wild-type cells (n = 130), roughly half the population contained 1-2 (dark blue) or 3-4 (light blue) Taz1-GFP foci across the cell cycle. Upon exposure to 1,6-hexanediol (n = 135), Taz1-GFP foci number increased, particularly during G2, indicating telomere de-clustering. In cells lacking Swi6 (n = 150), significant telomere de-clustering relative to wild-type cells was observed. * p < 0.05; **** p < 0.0001 by Fisher’s exact test.

Consistent with previous observations for HP1-α in mammalian cells^23^, the addition of 5% of the aliphatic alcohol 1,6-hexanediol for 15 minutes, which disrupts weak hydrophobic interactions, leads to both a rapid decrease in both the size and intensity of nuclear GFP-Swi6 foci as well as the accumulation of a diffuse pool of GFP-Swi6 in the cytoplasm (Figure S1). However, a pool of GFP-Swi6 is retained within the HC foci, which are expected to be 1,6-hexanediol-resistant due to direct binding via the Swi6 chromodomain to nucleosomes, particularly those containing methylated histone H3K9. Taken together, these observations support the existence of at least two populations of Swi6 molecules within HC foci: one directly bound to chromatin through the Swi6 chromodomain and the other driven by biomolecular condensation that is sensitive to 1,6-hexanediol, consistent with our prior single-molecule experiments demonstrating four distinct diffusive states of Swi6^44^. These observations suggest Swi6, as an ensemble, is an attractive candidate to impart viscoelastic properties to HC and, in turn, to the nucleus as a whole.

### Heterochromatin domains exhibit liquid-like properties *in vivo* that are tied to normal telomere clustering

As a role for biomolecular condensation in driving telomere clustering in cancer cells has been described previously^45,46^, we also suspected that liquid-like fusion could drive the coalescence of the six (replicated) heterochromatic telomeres, which are clustered together at the nuclear periphery in *S. pombe*^47^. When visualized by expressing a GFP fusion of Taz1, a shelterin complex subunit that specifically binds telomeric DNA^48^, approximately half of nuclei display one or two foci containing the six telomeres, with the other half displaying up to four foci (Figure 1C,D and Figure S2). This behavior is largely cell cycle-independent, as assessed using cell length as a proxy for cell cycle progression (Figure 1D). To further interrogate if telomere clustering relies on liquid-like condensation mediated by weak hydrophobic interactions, we treated cells with 1,6-hexanediol (5% for 30 minutes). This perturbation resulted in an increase in the number of Taz1-GFP foci per nucleus as the telomeres de-clustered (Figure 1D).

Similar to the 1,6-hexanediol treatment, we found that *swi6Δ* cells have an increase in the number of Taz1-GFP foci per nucleus (Figure 1D), indicating a clustering defect and a reliance on Swi6 for proper telomere coalescence. We note that even in this condition many cells display less than six discernible telomere foci, which we suspect occurs due to stochastic coincidence of telomeres in the nuclear volume or an inability to fully visualize the dimmer, de-clustered telomeres. Taken together, these observations suggest that Swi6 HC foci have a component of their behavior that is liquid-like and sensitive to treatment with 1,6-hexanediol. Furthermore, Swi6 contributes to the normal coalescence of telomere foci in fission yeast.

It is important to note that in fission yeast (as in mammalian cells), Swi6 shares its transcriptional silencing role in gene regulation with other chromodomain proteins, including Chp1 and Chp2, which are necessary for the proper establishment and balance of heterochromatic regions, respectively^49,50^. All three chromodomain proteins rely on the H3K9 methyltransferase Clr4 to be recruited to methylated histones of heterochromatic domains^49,51^. Their protein structures, however, differ from each other; for example, Chp1 lacks a chromoshadow domain and contains a much smaller N-terminal domain^49^ (Figure S3A). Under wild-type conditions, Chp1—like Swi6—binds to HC to form discrete foci (Figure S3B), and, also similar to Swi6, Chp1 is unable to bind and localize to heterochromatic foci in the absence of Clr4^49^ (Figure S3C,G). When H3K9 HC was allowed to spread in the absence of the H3K9me2/3 demethylase Epe1^52^, we observed that Chp1-GFP intensity increased at HC foci (Figure S3D-G), consistent with an enlargement of HC domains in Epe1-deficient cells. In the absence of Swi6, Chp1-GFP foci intensity also significantly increased (Figure S3E-G), suggesting competition of H3K9me2/3 binding between Swi6 and Chp1. With this, we can thus infer that the other chromodomain proteins, even at increased levels, are unable to replace Swi6’s role in telomere clustering in *swi6Δ* cells (Figure 1D), highlighting a unique feature of Swi6.

### The *swi6-sm1* allele disrupts the ability of HP1-α/Swi6 to promote the coalescence of heterochromatin domains

To determine the biochemical features of Swi6 tied to its ability to form and coalesce liquid-like HC domains, we turned to prior genetic analysis of Swi6 function. In particular, we interrogated the *swi6-sm1* allele, which carries two mutations in its N-terminal domain, rendering it unable to silence HC while leaving its H3K9me2/3-binding activity, dimerization, and cohesin interactions unaffected^53^ (Figure 2A). In previous studies of Swi6, defects in its N-terminal phosphorylation by casein kinase II also led to loss of transcriptional silencing^54^. Indeed, the D21G mutation encoded in the *swi6-sm1* allele disrupts the casein kinase II recognition site for phosphorylation of serine 18; this site is phosphorylated *in vivo* and *in vitro* by casein kinase II, and its targeted mutation disrupts Swi6’s ability to promote gene silencing^54^, thereby phenocopying the *swi6-sm1* allele. As N-terminal phosphorylation enhances oligomerization leading to phase separation for both HP1α and Swi6^22,55^, we hypothesized that the *swi6-sm1* allele could disrupt Swi6’s ability to undergo biomolecular condensation.

**Figure 2.**
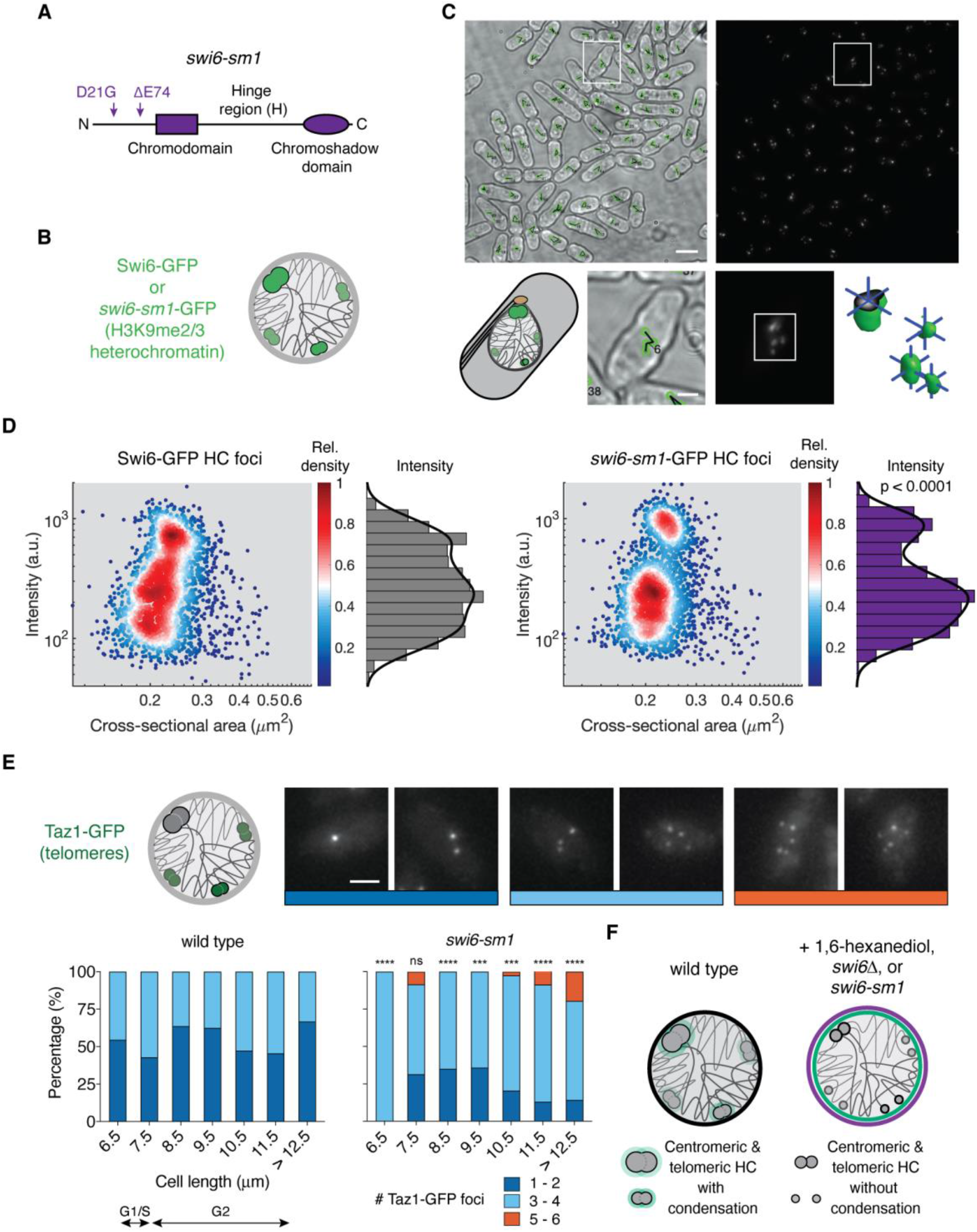
The *swi6-sm1* allele disrupts the ability of HP1-α/Swi6 to promote the coalescence of heterochromatin domains. (A) The *swi6-sm1* allele encodes two mutations in the N-terminal domain of Swi6. (B) Both wild-type Swi6-GFP and *swi6-sm1*-GFP localize to HC domains. (C) HC domain size and intensity measured using 3D domain reconstruction. Maximum intensity projections of transmitted light (upper left) and fluorescence images (upper right) of *S. pombe* cells expressing Swi6-GFP. Foci are grouped by cell (lower left) and individually reconstructed in three dimensions (lower right). Scale bars, 6 μm and 2 μm. (D) The *swi6-sm1* allele leads to a depletion of intermediate scale and intense heterochromatic domains relative to wild-type Swi6-GFP. Foci intensity versus cross-sectional area density scatter plots for Swi6-GFP (n = 2500 foci) and *swi6-sm1*-GFP (n = 2402 foci) labeled HC domains. **** p < 0.0001 by Kolmogorov-Smirnov test after calculation of cumulative distributions. (E) The *swi6-sm1* allele leads to telomere de-clustering in a manner similar to 1,6-hexanediol exposure and *swi6Δ*. Data expressed as (and wild-type data replotted from) in Figure 1D. In cells expressing *swi6-sm1* (n = 145), the de-clustering effect recapitulates that seen with 1,6-hexanediol-treated cells or cells lacking Swi6. *** p < 0.001; **** p < 0.0001 by Fisher’s exact test. (F) Summary of HC domain phenotypes. In wild-type cells, HC domains form discrete clusters. Treatment with 1,6-hexanediol, deletion of Swi6, or expression of the N-terminal mutant *swi6-sm1* causes HC domains to decluster.

Like WT Swi6-GFP, *swi6-sm1*-GFP accumulates in foci within the nucleus close to the nuclear periphery (Figure 2). To begin to probe if and how the *swi6-sm1* mutation alters the coalescence of HC domains, we compared the size, intensity, and number of Swi6-GFP and *swi6-sm1*-GFP foci within individual nuclei (Figure 2B). To quantitatively compare foci size and intensity, we employed an image analysis approach we previously established that reconstructs HC foci from image z-stacks with sub-pixel resolution^56^ (Figure 2C). Comparing each foci’s intensity to its cross-sectional area in an ensemble scatter plot revealed divergent populations in Swi6-GFP versus *swi6-sm1*-GFP-labeled HC (Figure 2D and Figure S4). In wild-type cells, Swi6-GFP HC domains comprise a nearly uniform distribution of intensities ranging from bright (centromeric HC) to dim (telomeric and other HC foci). Cells expressing *swi6-sm1*-GFP, however, display a depletion of intermediately intense HC foci, indicative of non-centromeric HC domain de-clustering (p < 0.0001). This depleted population is most likely to represent intact, condensed telomeric domains, indicating that *swi6-sm1* affects proper telomere coalescence. To support this, expression of the *swi6-sm1* allele with Taz1-GFP shows disruption of normal telomere clustering similar to complete loss of Swi6 or treatment with 1,6-hexanediol (Figure 2E and Figure 1D). To the extent that the condensation of Swi6 facilitates proper telomere clustering, we conclude that the *swi6-sm1* allele could specifically disrupt aspects of Swi6 function that underlie its ability to form condensates (Figure 2F).

### The *swi6-sm1* allele shows a depletion of mobile states despite normal chromatin and H3K9me2/3 binding *in vivo*

Previously, we used a single-molecule approach to track the dynamics of Swi6 in the fission yeast nucleus^44^. We fused Swi6 to the photoactivatable fusion protein PAmCherry and captured the distinct biophysical states associated with its motion within its native chromatin context (Figure 3A,B). Through mutational analysis, we defined how each biophysical state of Swi6 is associated with a unique biochemical or structural property that influences how Swi6 behaves at sites of H3K9 methylation^44^. Our analysis revealed four states of WT PAmCherry-Swi6 molecules with different diffusion coefficients (Figure 3C, crosses) and transition probabilities between the states (Figure 3D). The least mobile state (termed α, blue) comprises molecules bound to H3K9me2/3 (weight fraction, *π* ∼25%), the intermediate states indicate molecules associated with unmethylated H3K9 nucleosomes (termed β, orange; *π* ∼33%) or nucleic acids (termed γ, yellow; *π* ∼15%), and the remaining, most mobile state is comprised of unbound molecules (termed δ, purple; *π* ∼27%) (Figure 3C)^44^. Note that this observed high-mobility state of Swi6 displays a ∼40-fold slower diffusion coefficient (D = 0.51 μm^2^/s) than what has been reported for similar-size GFP-tagged HP1 monomers and dimers in euchromatic regions where it is expected to be relatively fully unbound/free^57^. Thus, the δ state is slowed significantly compared to truly freely diffusing molecules, consistent with it representing unbound but restrained movement in the mobile phase of the condensate at HC domains. Indeed, the rapid diffusion of completely free monomers and dimers would not detectable within the acquisition parameters optimized for our PAmCherry-Swi6 experiments.

**Figure 3.**
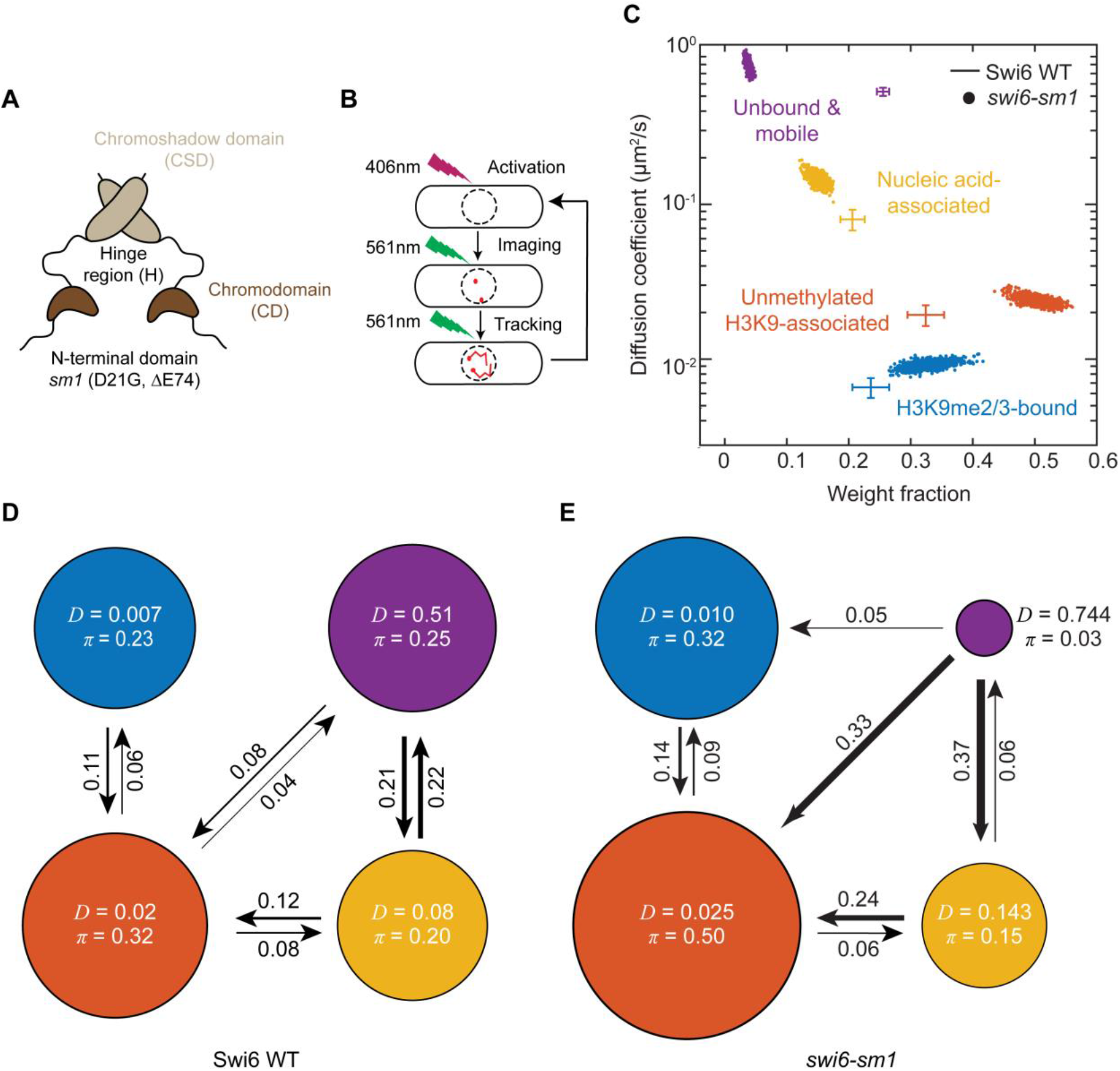
The *swi6-sm1* allele shows a depletion of mobile states despite normal chromatin and H3K9me2/3 binding *in vivo*. (A) Schematic representation of Swi6. Each domain has a distinct biochemical property. NT: unstructured N-terminus subject to phosphorylation that regulates Swi6 condensation *in vitro* (*swi6-sm1* has two mutations as indicated); CD: H3K9me binding; H: nucleic acid binding; CSD: protein oligomerization. (B) Single-molecule tracking photoactivation localization microscopy (spt-PALM) of wild-type Swi6 and *swi6-sm1* in the *S. pombe* nucleus. 1-3 PAmCherry-Swi6 molecules/nucleus are photoactivated (406 nm, 1.50-4.50 W/cm2, 200 ms), then imaged and tracked until photobleaching (561 nm, 71.00 W/cm2, 25 frames/sec). The cycle is repeated 10-20 times/cell. (C) SMAUG identifies four distinct mobility states, (α blue, β orange, γ yellow, and δ purple) for PAmCherry-Swi6 and PAmCherry-*swi6-sm1*. Each point represents the average single-molecule diffusion coefficient, D, versus weight fraction of Swi6/*swi6-sm1* molecules in that state at a saved iteration of the Bayesian algorithm after convergence. Dataset, PAmCherry*-swi6-sm1*: 29,657 steps from 3,744 trajectories and 97 cells. Crossbars: comparison to PAmCherry-Swi6 results from reference ^44^ with 10,095 steps from 1,491 trajectories and 36 cells. (D) Average probabilities (arrows) of a PAmCherry-Swi6 molecule transitioning between the mobility states (circles) from reference^44^. Each circle area is proportional to the weight fraction, π; D is in µm^2^/s. Low-significance transition probabilities below 0.04 are excluded. (E) Average probabilities (arrows) of a PAmCherry-*swi6-sm1* molecule transitioning between the mobility states (circles) as in (D) (see also Table S1).

In light of the quantitative defects observed in the HC domains formed in the presence of the *swi6-sm1* allele (Figure 2), we sought to characterize how *swi6-sm1* mutations affect the distribution of mobility states and thus infer how its two N-terminal mutations influence Swi6 dynamics. Overall, PAmCherry-*swi6-sm1* molecules move more slowly compared to PAmCherry-Swi6 as noted by the ∼30% increase in the fraction of molecules in the H3K9me and chromatin bound states (α and β states) (Figure 3C, scatter dots). Using Single-Molecule Analysis by Unsupervised Gibbs sampling (SMAUG) analysis^58^, we distinguished four mobility states associated with PAmCherry-*swi6-sm1*; the same number as previously observed for wild-type PAmCherry-Swi6 (Figure 3C,E). The *swi6-sm1* mutations, however, led to significant changes in the distributions of molecules within each state (Figure 3C) and the transitions between them (Figure 3E and Table S1). Of note, the weight fraction, π, of the mobile (δ) Swi6 population, previously associated with unbound Swi6 molecules, decreased significantly from *π* = 27% to only 3% of all molecules. Concomitant with this decrease, there is an increase in the H3K9me2/3-bound (α) state weight fraction from *π* = 25% to 32% and a substantial increase in the occupancy of the nucleosome-bound (β) state from *π* = 33% to 50%. We also analyzed the transition probabilities between the different mobility states and observed that a major difference between WT Swi6 and *swi6-sm1* is a 4-fold increase in the transition probability between the δ and β states, suggesting that the small pool of free *swi6-sm1* molecules is likely to associate with chromatin (H3K9me0 or H3K9me2/3). Taken together, these results confirm that PAmCherry-*swi6-sm1* is fully competent to associate with chromatin and is, in fact, more likely to remain bound to nucleosomes compared to wild-type Swi6. Importantly, the increase in the percentage of chromatin-bound PAmCherry-*swi6-sm1* molecules leads to a profound depletion of the mobile PAmCherry-*swi6-sm1* pool and a substantial reduction in the likelihood of returning to the unbound state from the chromatin-bound state, a feature that potentially correlates with the fluidity of liquid-like Swi6 condensates^59^.

### The *swi6-sm1* mutations disrupt the ability of HP1-α/Swi6 to form a condensed, mobile phase around H3K9me2/3 chromatin

To further interrogate how the *swi6-sm1* mutations affect its global condensation at HC domains, we imaged diploid cells expressing either (1) one copy of WT Swi6-GFP and one copy of WT Swi6-mCherry (Figure 4A) or (2) one copy of WT Swi6-GFP and one copy of *swi6-sm1*-mCherry (Figure 4B). We hypothesized that the depletion of the condensed, mobile fraction of *swi6-sm1* molecules as revealed by our single molecule experiments would manifest as a decrease in the total number of Swi6 molecules at HC foci alongside an otherwise normal pool of relatively immobile *swi6-sm1* molecules. Indeed, while both Swi6-GFP and *swi6-sm1*-mCherry co-localize within HC domains as expected due to intact H3K9me2/3 and dimerization properties (Figure 3), WT Swi6 accumulates to a greater extent, forming brighter foci than the mutant *swi6-sm1* (Figure 4B) when compared to images of the WT/WT diploid, which we used to normalize the scale between the co-localized GFP and mCherry foci (Figure 4C and Figure S5). When averaged over thousands of HC foci, the mean WT Swi6-GFP foci intensity profiles were similarly bright across the two diploid strains, while the mean *swi6-sm1*-mCherry foci intensity was reduced by ∼25% compared to the WT Swi6-mCherry foci (Figure 4D). Comparison of the brightest, intermediate, and dimmest foci within each diploid nucleus revealed similar reductions in *swi6-sm1* concentration, suggesting the mutant allele affects Swi6’s condensation properties across all sizes of HC domains, with bright foci corresponding to centromeric HC and intermediate to dim foci representing telomeric and other smaller HC domains (Figure 4E). Our findings suggest that one pool of Swi6 is bound directly to HC (intact in the *swi6-sm1* allele) while a second pool is localized to and concentrated at HC domains via its condensation (disrupted in the *swi6-sm1* allele). Taken together, our data argue that the *swi6-sm1* allele represents a separation-of-function mutant with a specific defect in Swi6 condensation.

**Figure 4.**
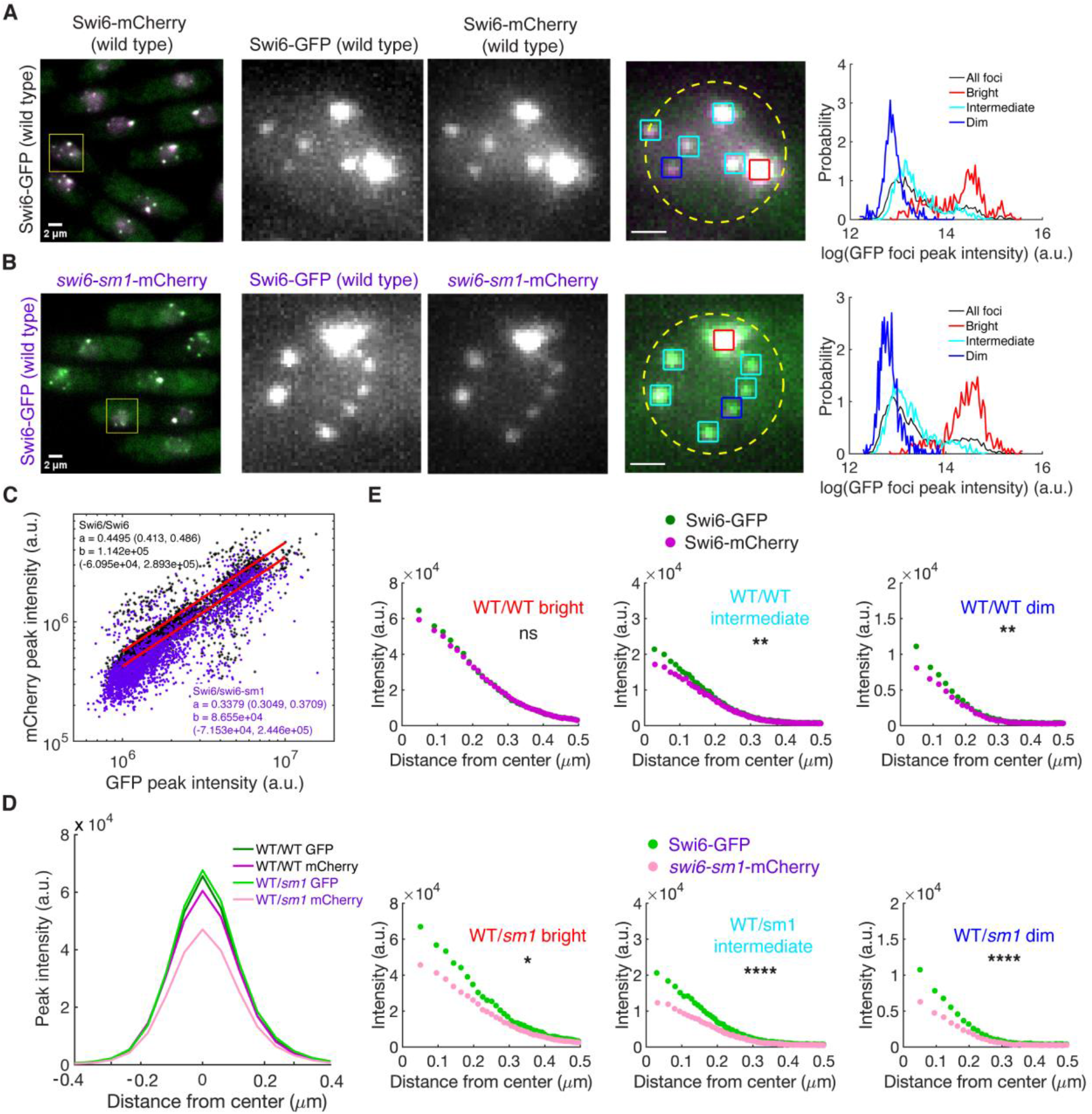
The *swi6-sm1* mutations disrupt the ability of HP1-α/Swi6 to form a condensed, mobile phase around H3K9me2/3 chromatin. (A) Maximum intensity projections of fluorescence images of diploid *S. pombe* cells expressing Swi6-GFP/Swi6-mCherry. Inset shows a representative nucleus with detected Swi6 domains marked by intensity as bright (red, the brightest domain in the nucleus), dim (blue, the dimmest domain), and intermediate (cyan, all the remaining domains). Right panel shows distribution of “bright”, “intermediate”, “dim” domains for combined domains across all cells with ≥ 3 foci. For display the intensity between GFP and mCherry was normalized (calibration from (C)). Scale bar, 2 μm, inset, 1 μm. (B) Same as (A) but for cells expressing Swi6-GFP/*swi6-sm1*-mCherry and using the same calibration between channels as in (A). (C) Scatter plot of Swi6 foci intensities in diploid cells from nuclei with ≥ 3 foci, shown as mCherry signal versus GFP signal for Swi6-GFP/Swi6-mCherry cells (black, n = 584 nuclei; 2,648 foci) and Swi6-GFP/*swi6-sm1*-mCherry cells (purple, n = 538 nuclei; 2,327 foci). Red lines are best fit of a linear model y(x) = a*x+b, with a = 0.4495, b = 1.142 x 10^5^ and a = 0.3379, b = 8.655 x 10^4^, respectively. Values in parentheses are 95% confidence intervals. Coefficient a = 0.4495 was used to rescale mCherry channel fluorescence in (A) and (B) (see also Figure S5). (D) Average intensity profile for GFP and mCherry fluorescence for Swi6 domains after mCherry signal rescaling using calibration from (C). (E) Pixel intensity as a function of distance from the center of Swi6 domain (determined in GFP channel) for GFP and mCherry signal for “bright”, “intermediate”, and “dim” domains. Upper panels: Swi6-GFP/Swi6-mCherry cells; bottom panels: Swi6-GFP/*swi6-sm1*-mCherry cells. * p < 0.05; ** p < 0.01; **** p < 0.0001 by two-sample Kolmogorov-Smirnov test.

### H3K9me2/3 and the condensed, mobile pool of Swi6 are required to impart nuclear stiffness *in vitro*

As we previously demonstrated that peripherally-tethered chromatin at the inner nuclear membrane is a source of mechanical support for the nucleus^12^, we sought to evaluate the mechanical contribution of Swi6, and specifically its ability to undergo condensation, on nuclear mechanics. To directly interrogate how fission yeast nuclei respond to explicit magnitudes, time scales, and compression/tension force regimes, we employed an *in vitro* force spectroscopy approach on isolated nuclei that we developed previously^12^ to directly measure how altering the extent of H3K9 methylation or loss of HP1-α/Swi6 condensation impacts nuclear stiffness.

Nuclei expressing a GFP fusion of the nucleoporin Cut11 were isolated and used to perform single-nuclei force spectroscopy measurements using optical tweezers (Figure 5). To directly measure its mechanics, a nucleus is sandwiched between two silica beads, one a larger bead adherent to a glass slide with the other, smaller bead trapped in the optical trap (Figure 5A,B). As the larger bead is moved by a piezo stage (Δ*x*_piezostage_), the displacement of the smaller bead in the trap is measured (Δ*x*_bead_); the difference in x-displacement is the amount by which the nucleus is either stretched or compressed (Δ*x*_nucleus_). Each oscillation yields force and displacement measurements that can be plotted with respect to each other (Figure 5C). The oscillation data display a linear relationship (black line); the slope (k) of which is equivalent to the spring constant, or effective stiffness, of the individual nucleus. Each nucleus is subjected to multiple rounds of tension and compression at varying time scales, with all oscillations performed over a range of biologically-relevant frequencies: 0.01 Hz – 2 Hz (Figure S6).

**Figure 5.**
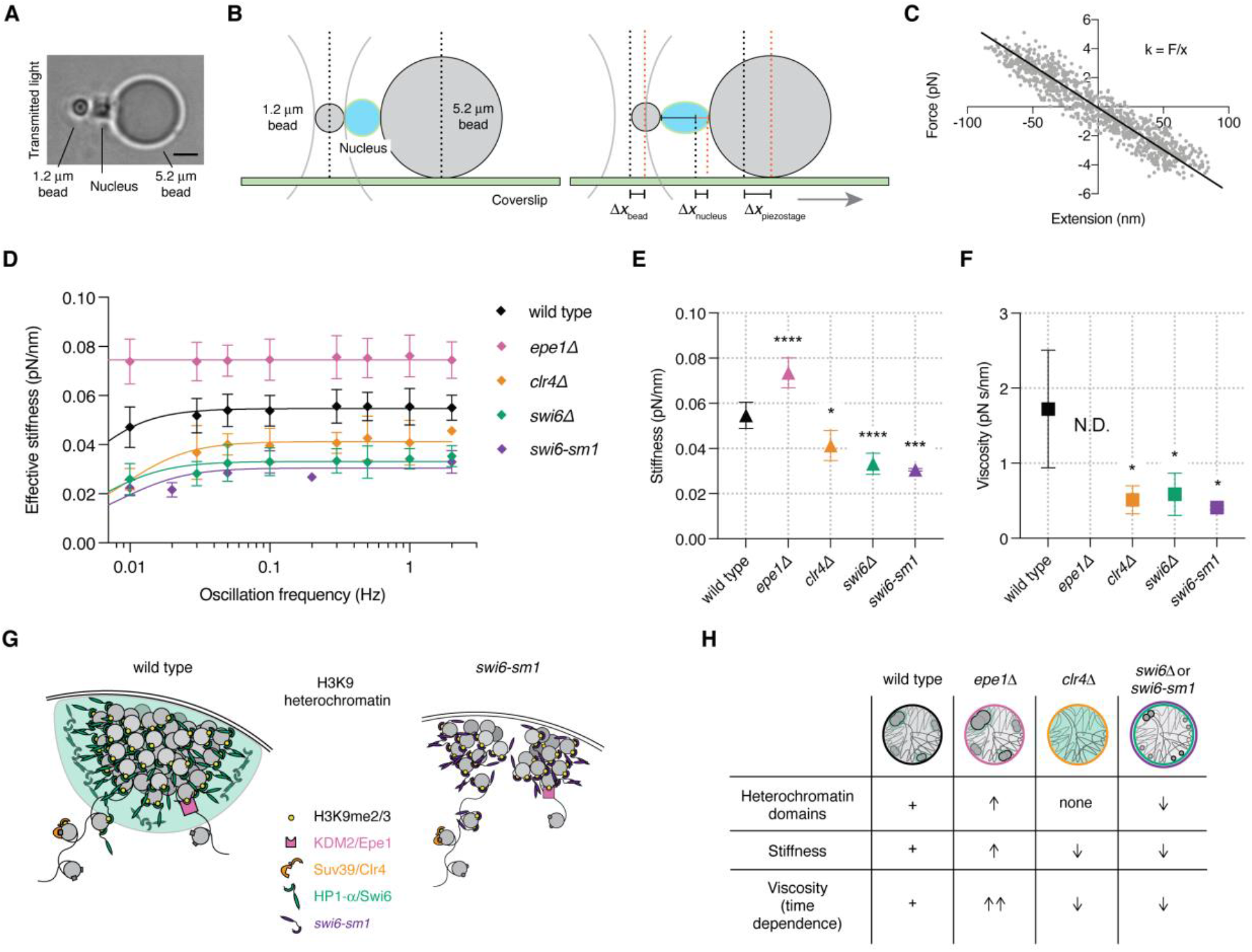
H3K9me2/3 and the condensed, mobile pool of Swi6 are required to impart nuclear stiffness as assessed *in vitro*. (A) Transmitted light image showing the force spectroscopy configuration as viewed from above. Scale bar, 2 μm. (B) Schematic representation of the force spectroscopy assay. (C) Representative force versus extension curve of an isolated wild type nucleus. Each data point (gray) represents the force applied to the nucleus and the resulting displacement over one oscillation (extension and compression). The data display a linear relationship (black line); the slope (k) is equivalent to the spring constant, or effective stiffness, of the individual nucleus. (D) Effective stiffnesses of wild type nuclei (black, n = 8 nuclei; 63 oscillations), KDM2/Epe1-deleted nuclei (pink, n = 18 nuclei; 172 oscillations), Suv39/Clr4-deleted nuclei (orange, n = 3 nuclei; 22 oscillations), HP1-α/Swi6-deleted nuclei (green, n = 5 nuclei; 40 oscillations), and *swi6-sm1-*expressing nuclei (purple, n = 3 nuclei; 29 oscillations) over a range of oscillation frequencies (0.01 – 2 Hz). Data are represented as mean +/- SD of the raw oscillation data at each oscillation frequency. (E) Nuclear stiffnesses extracted from viscoelastic fit of data in (D). Data are represented as mean +/- SD of the fitted values. (F) Nuclear viscosity extracted from viscoelastic fit of data in (D). Data are represented as mean +/- SD of the fitted values. N.D., not determined. (G) Models of H3K9me HC with (left) and without (right) Swi6 condensation. Nucleosomes (gray) are methylated on histone H3K9 (yellow) by the methyltransferase Suv39/Clr4 (orange). Upon methylation, the HC protein Swi6 (green) binds H3K9me2/3, crosslinking adjacent nucleosomes to form compact HC. Condensation of additional, unbound Swi6 molecules form an extended phase nucleated at these H3K9me2/3 chromatin regions (represented by green cloud). The demethylase KDM2/Epe1 (pink) reverses this process, countering HC formation by promoting H3K9me2/3 demethylation. *swi6-sm1* (purple) binds to, crosslinks, and compacts H3K9me2/3 nucleosomes normally but disrupts the ability to condense and form this mobile phase. (H) Schematic table representing the mechanical effects of altering H3K9me and binding proteins. * p < 0.05; ** p < 0.01; *** p < 0.001; *** p < 0.0001 by ordinary one-way ANOVA with Dunnett’s correction for multiple comparisons.

The response to compression and tension forces revealed a pronounced mechanical defect in nuclei purified from a strain in which the Suv39/Clr4 H3K9 methylase was deleted, which is necessary for H3K9me2/3 HC domain formation and therefore lies upstream of Swi6. These nuclei displayed a ∼50% decrease in effective stiffness over the range of oscillation frequencies (Figure 5D,E). Interestingly, loss of the presumed H3K9me2/3 demethylase KDM2/Epe1, which leads to a moderate expansion of HC domains^52^ (Figure S3D,F), leads to a ∼30% increase in nuclear stiffness (Figure 5D,E), suggesting that the extent of H3K9 methylation can dictate the mechanical response of the nucleus to deformation.

We next tested whether Swi6 contributes to the mechanism by which the extent of H3K9 methylation impacts nuclear stiffness. Strikingly, *swi6Δ* nuclei also had a ∼50% decrease in stiffness, phenocopying *clr4Δ* nuclei (Figure 5D) and suggesting that Swi6 is integral to the mechanism by which H3K9me imparts nuclear stiffness. To investigate the aspect(s) of Swi6 function that underlie its mechanical contribution, we next evaluated the mechanical behavior of *swi6-sm1* nuclei. To our surprise, although *swi6-sm1* appears to be entirely competent for its direct biochemical association with H3K9me2/3 nucleosomes (Figure 3), these nuclei are mechanically deficient to the same extent as *swi6Δ* nuclei (Figure 5D). This finding suggests that it is the mobile molecules of Swi6 condensed at HC domains (not the molecules bound directly to H3K9me or nucleosomes) that play a critical role in the contribution of Swi6 to nuclear stiffness.

We previously observed that fission yeast nuclei display a viscoelastic response^12^. Repeating oscillations at different frequencies can reveal the time-dependent (viscous) component of the nuclear response (see Figure S6). At longer oscillation frequencies (100 s or 0.01 Hz), we observed a characteristic reduction in effective stiffness—a non-linear mechanical response—in wild type, *clr4Δ*, *swi6Δ*, and *swi6-sm1* nuclei, indicating that these systems are not perfect springs but instead combinations of an elastic spring and a viscous dashpot, or dampener. To extract the individual mechanical properties of nuclear stiffness and viscosity, we performed non-linear regression applying the Maxwell viscoelastic model to our frequency series of effective stiffnesses (Figure 5D, solid lines). From the viscoelastic fits in (D), the nuclear stiffness was extracted for each genotype. As compared to the mean stiffness of wild-type nuclei (0.055 pN/nm), *epe1Δ* nuclei were stiffer (0.073 pN/nm), and *clr4Δ*, *swi6Δ,* and *swi6-sm1* nuclei were softer (0.041 pN/nm, 0.033 pN/nm, and 0.030 pN/nm respectively) (Figure 5E).

Relative to the viscosity of wild type nuclei (1.72 pN s/nm), *clr4Δ*, *swi6Δ,* and *swi6-sm1* nuclei were also significantly less viscous (0.51 pN s/nm, 0.58 pN s/nm, and 0.41 pN s/nm, respectively) (Figure 5F). In contrast, loss of Epe1 also led to an increase in nuclear viscosity to the extent that we were unable to be determine its value from the timescales sampled with our oscillation assay (note the complete time-independence of the measured stiffness in Figure 5D). Thus, despite a rather subtle effect on the recruitment of *swi6-sm1* to HC domains via its intact H3K9me2/3 binding and dimerization activities (Figures 3 and 4), the loss of its ability to undergo condensation and/or to drive coalescence of HC domains is critical to its mechanical contribution to nuclear mechanics (Figure 5G,H).

### Loss of HP1-α/Swi6 condensation increases microtubule-driven nuclear deformations *in vivo*

To assess how loss of HP1-α/Swi6 or abrogation of its condensation affects the nuclear response to MT dynamics, which primarily exert force through the SPB on the centromeric HC domain, we monitored nuclear envelope (NE) deformations *in vivo* (Figure 6A). To evaluate these nuclear deformations with precise spatial and temporal resolution, the nuclear envelope was labeled by expression of the nucleoporin Cut11-GFP and visualized in all four-dimensions (4D: x, y, z, and time) by live-cell fluorescence microscopy (Figure 6B). We next employed a three-dimensional (3D) surface reconstruction software we previously developed^12,56^ applied to series of fluorescent z-stacks with sub-pixel resolution to capture intrinsic MT-driven deformations (Figure 6C). As expected, the largest NE deformations manifest at points subjected to maximum shear forces as MTs drive SPB oscillations, which can be visualized as local maxima on the nuclear surface (Figure 6B,C,D, left column); these maxima are lost upon MT depolymerization with the drug carbendazim (MBC) (Figure 6D, right column, and E, dashed lines). When averaged over many nuclei, the largest NE deformations occur at ∼45 and 135 degrees with respect to the long axis of the cell (Figure 6E, right panel, and Figure S7)—the points at which the SPB is maximally translated due to the shearing force from the MTs (Figure 6A).

**Figure 6.**
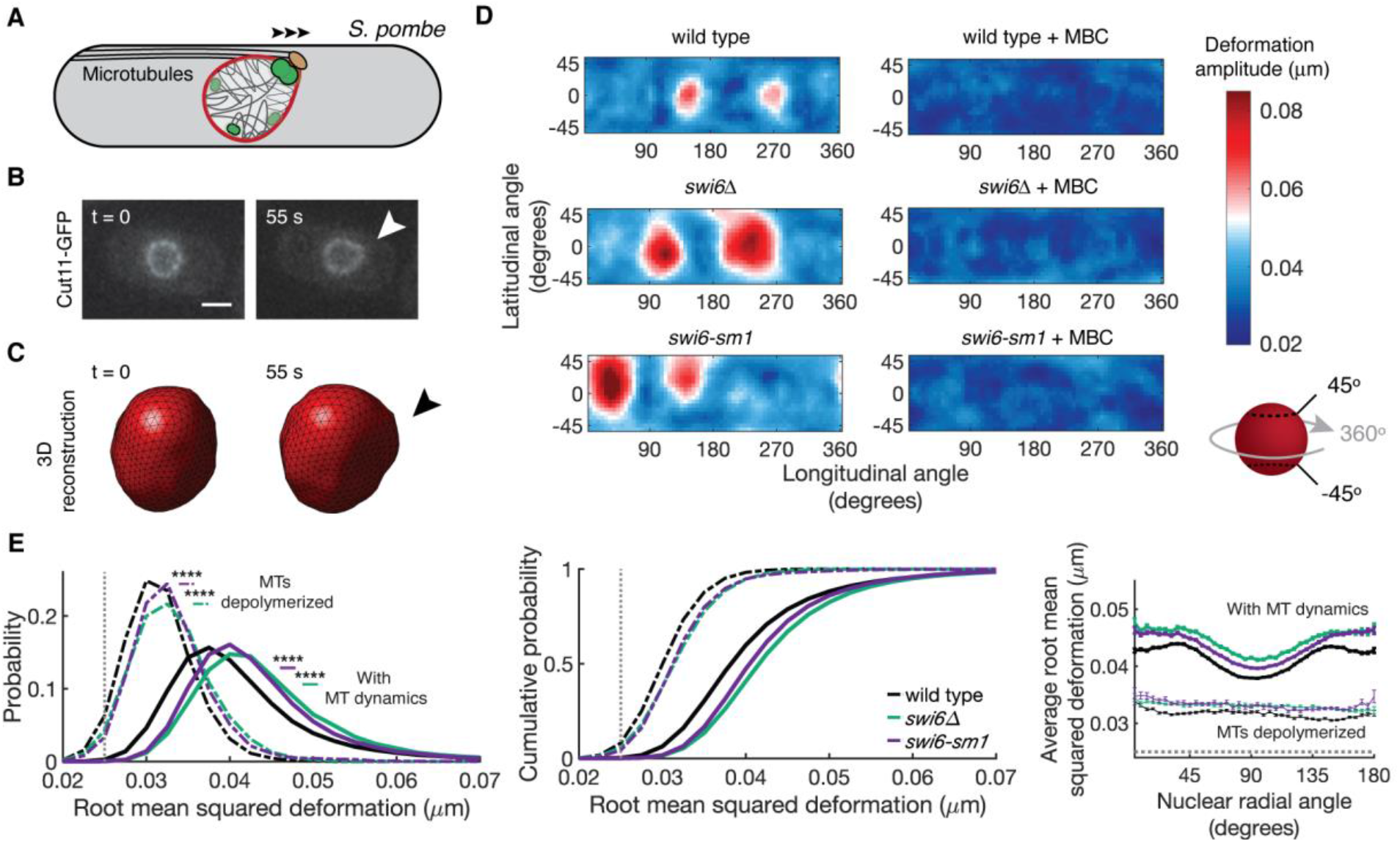
Loss of HP1-α/Swi6 or its condensation increases microtubule-driven nuclear deformations *in vivo*. (A) In *S. pombe*, shearing forces acting upon the nuclear envelope (red) arise from polymerizing microtubules (MTs, black) at the cell tips that drive oscillation of the associated spindle pole body (SPB, tan) and are transmitted to the centromeric HC (large green domain). (B) Fluorescence images of a wild. type *S. pombe* cell expressing Cut11-GFP demonstrating a typical nuclear deformation (arrowhead). Scale bar, 2 μm. (C) 3D reconstructions of the nucleus. At t = 0 (left), the nucleus is circular; ∼55 seconds later (right), the same nucleus undergoes a MT-dependent deformation (arrowhead). (D) Representative single nuclear surface deformations with (left column) and without (right column) MT dynamics shown as topological heat maps with respect to its latitudinal angle (points above and below the nuclear “equator”)—a band between +45 and -45 degrees—and its longitudinal angle (points around the nucleus) between 0 and 360 degrees (see also Figure S7). (E) Root mean squared deformation size across all nuclei observed (left and middle) and all nuclear radial angles (right): wild type (black, n = 760 nuclei; 12,556 deformations), *swi6Δ* (green, n = 218 nuclei; 4,421 deformations), *swi6-sm1* (purple, n = 371 nuclei; 6,583 deformations). MBC-treated nuclei shown as dashed lines: wild type (black, n = 58 nuclei; 561 deformations), *swi6Δ* (green; n = 69 nuclei; 976 deformations), *swi6-sm1* (purple, n = 36 nuclei; 534 deformations). Data are represented as mean (+/- SEM in right plots in (E)). **** p < 0.0001 by Mann-Whitney U test.

Similar to the mechanical defect seen in our *in vitro* experiments, cells lacking Swi6 displayed larger deformations at all angles as compared to wild-type cells, as reflected in the root mean squared deformation size in both the presence and absence of MT dynamics (Figure 6E, left and middle, p < 0.0001). Averaging over hundreds of nuclei and thousands of individual NE deformations, the locations of the deformations in *swi6Δ* cells mirrored the wild-type profile, but were shifted towards larger deformations (Figure 6E, right). This suggests that loss of Swi6 leaves the geometry of forces exerted on the nucleus intact while reducing its ability to resist MT forces (in contrast to loss of Suv39/Clr4, which cannot be interpreted because this genetic background disrupts the characteristic MT-induced nuclear deformations, see Figure S8A,B).

Cells lacking Swi6 have a defect in cohesin loading at the centromeres^60^, which are subjected to tangential MT forces *in vivo* (Figure 1). In contrast, the *swi6-sm1* allele retains the ability to support normal centromeric cohesion^41^ and to associate with H3K9me2/3 nucleosomes (Figure 3), but depletes the mobile phase of Swi6 molecules and limits HC condensation of smaller domains (Figures 2-4). This allele therefore provides an opportunity to investigate the Swi6 functions(s) that contribute most to the response of the nucleus at the centromeric HC-MT interface. We observe that *swi6-sm1* cells largely recapitulated the increase in nuclear deformability observed in *swi6Δ* cells, albeit to a slightly lesser extent (Figure 6E, p < 0.0001 as compared to wild type). Taken together, these observations suggest that defects in Swi6 condensation at HC domains plays a more dominant role in the global, emergent viscoelasticity of the nucleus as a system measured *in vitro*, while there is a clear but less quantitative effect on local deformations in reponse to forces exerted on the centromeric HC-MT interface *in vivo*.

### Expansion of HC or crosslinking of H3K9me2/3 nucleosomes by Swi6 are insufficient to impart nuclear stiffness in the absence of a mobile, condensed pool of Swi6

Cells harboring the *swi6-sm1* allele have HC domain clustering defects (Figure 2), reduced Swi6 concentration at HC sites (Figure 4), and ultimately loss of global nuclear stiffness (Figures 5 and 6), all features we couple to the depletion of the mobile pool of Swi6 molecules even as the enhanced association of *swi6-sm1* molecules bound to (and cross-linking of) nucleosomes (Figure 3) is insufficient to retain these features. To further probe this model in the context of WT Swi6, we sought to understand how expansion of HC domains affects Swi6 binding states, and, in turn, how this expansion affects nuclear mechanics. By single molecule tracking, we found that loss of H3K9me2/3 demethylation activity (by deletion of the H3K9me2/3 demethylase, KDM2/Epe1) does not significantly alter the relative populations of bound versus unbound Swi6 as compared to WT cells (Figure 7A, left). However, combined loss of Epe1 and Mst2, a histone acetylase that enhances HC spreading when deleted^52^, leads to WT Swi6 being nearly entirely chromatin-bound, while the mobile pool normally found in the condensate is depleted^44^ (Figure 7A, right), recapitulating the phenotype seen in *swi6-sm1* cells (Figure 3C). We then tested whether HC spreading affects the *in vivo* nuclear force response. In this *in vivo* assay, which measured the local response to MT dynamics, we observed no significant difference in NE deformations in Epe1-deficient nuclei (Figure 7B and Figure S8C,D). Strikingly, however, we found that *epe1Δmst2Δ* cells displayed much larger NE deformations than WT or *epe1Δ* cells (Figure 7B), showing a paradoxical increase in nuclear deformability when HC spreads. We conclude that enhanced levels of H3K9me2/3 substantial enough to drive depletion of the mobile pool of Swi6 from the condensates compromises rather than reinforces nuclear mechanics.

**Figure 7.**
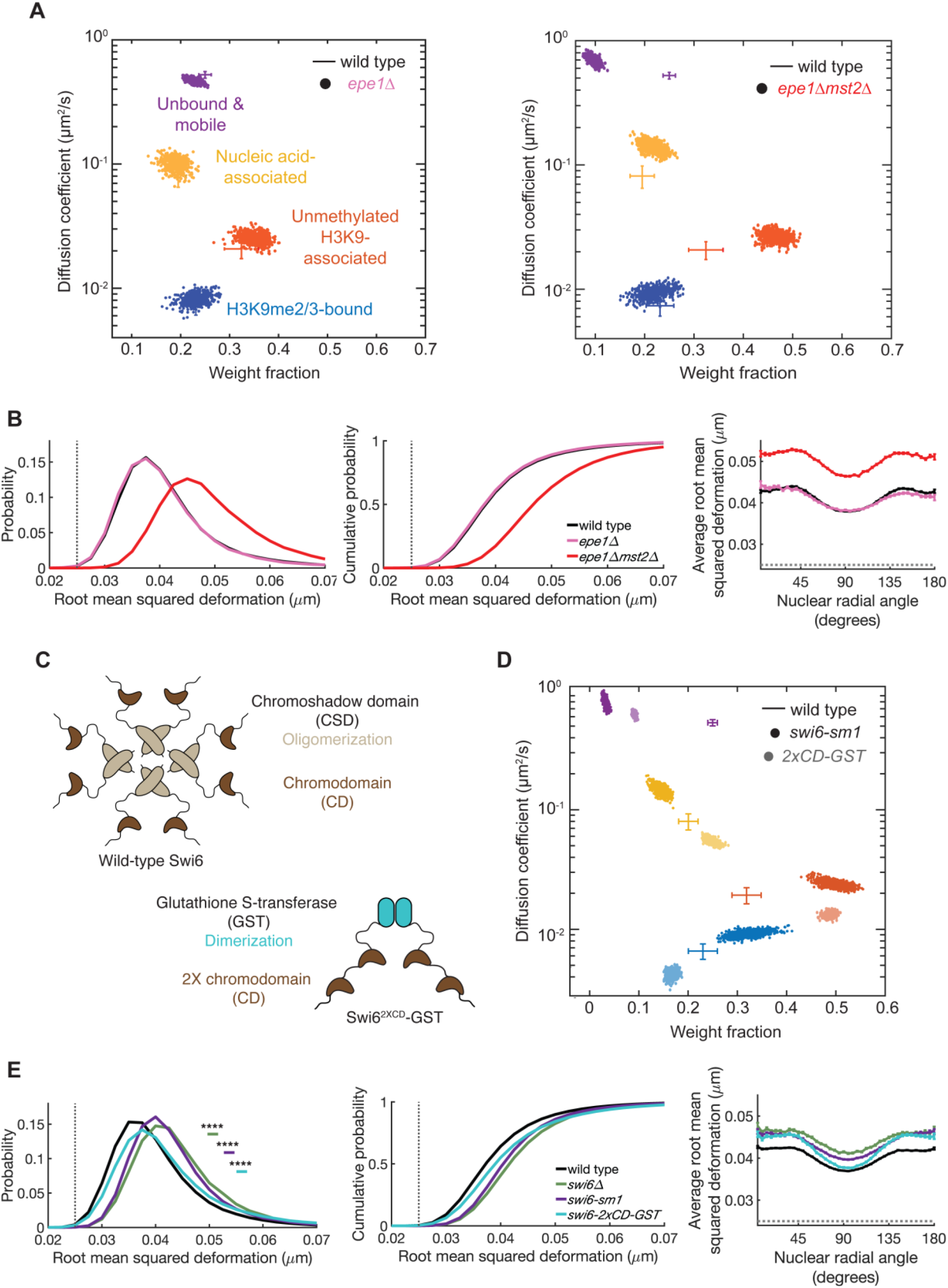
Expansion of HC or crosslinking of H3K9me2/3 nucleosomes by Swi6 are insufficient to impart nuclear stiffness in the absence of a mobile, condensed pool. (A) Depletion of both Epe1 and Mst2 (but not Epe1 alone), which drives heterochromatin spreading, leads to depletion of the mobile pool PA-mCherry-Swi6. SMAUG analysis and plots as in Figure 3C. Individual points represent the cells lacking Epe1 (left) or Epe1 and Mst2 (right) correspond to the indicated mobility states with the wild-type genotype shown by the cross-hatches (data from reference ^44^). (B) Expanding HC domains drives larger nuclear deformations. Root mean squared deformation size across all nuclei observed (left and middle) and all nuclear radial angles (right): wild type (black, n = 760 nuclei; 12,556 deformations, replotted from Figure 6E), *epe1Δ* (pink, n = 215 nuclei; 3,610 deformations), *epe1Δmst2Δ* (red, n = 361 nuclei; 8,419 deformations). Data are represented as mean (+/- SEM in right plots in (B)). (C) Diagram of wild-type Swi6 that assembles by oligomerization mediated by the chromoshadow domains (top) and an engineered H3K9me nucleosome crosslinking construct, PA-mCherry*-swi6-2xCD-GST* (developed in reference ^44^). (D) The PA-mCherry*-swi6-2xCD-GST* construct (light colors, data from reference ^44^; 42,382 steps from 5,182 trajectories and 140 cells), which lacks the domains mediating Swi6 condensation, largely phenocopies the diffusive states displayed by PAmCherry-*swi6-sm1* (dark colors). Wild-type PAmCherry-Swi6 (crosshatches) and PAmCherry-*swi6-sm1* data are re-plotted from Figure 3C (see also Table S1). (E) Expression of the *swi6-2xCD-*GST construct leads to larger nuclear deformations. Root mean squared deformation size across all nuclei observed (left and middle) and all nuclear radial angles (right): wild type with PAmCherry-Swi6 (black, n = 1,502 nuclei; 27,059 deformations), *swi6Δ* (green, n = 218 nuclei; 4,421 deformations; replotted from Figure 6E), *swi6-sm1* (purple, n = 371 nuclei; 6,583 deformations; replotted from Figure 6E), and *swi6-2xCD-GST* (teal, n = 329 nuclei; 6,055 deformations). Data are represented as mean (+/- SEM in right plots in (E)). **** p < 0.0001 by Mann-Whitney U test.

To further challenge this interpretation, we tested the sufficiency of an engineered H3K9me2/3-containing nucleosome crosslinker comprised of a GST moiety (which inherently drives dimerization^44,61^) fused to one or two units of the Swi6 chromodomain (CD) to influence nuclear deformability in cells lacking WT Swi6 (Figure 7C). These heterologous constructs, which lack the domains implicated in Swi6 condensation^44,62^, show a similar depletion of the mobile/unbound state and are found predominantly bound to chromatin when examined by single molecule tracking (Figure 7D). Concomitantly, we observed increased nuclear deformability upon expression of *swi6-2xCD-GST* (Figure 7E), highlighting again that driving H3K9me2/3 nucleosomes into dense arrays cannot support nuclear mechanics and indeed poisons mechanical robustness of the nucleus in the absence of WT Swi6, which otherwise would be condensed at HC domains. Thus, we conclude that Swi6 plays a key role in defining nuclear stiffness not by its HC-binding and crosslinking abilities but rather by its ability to undergo condensation into HC-associated droplets.

## DISCUSSION

In an effort to understand the contribution of heterochromatin to nuclear mechanics, we combined the use of specific H3K9me-associated genetic variants with a complementary array of *in vivo* and *in vitro* assays aimed at elucidating its biophysical mechanism. Our results support a model in which the mobile, condensed phase of the HC protein HP1-α/Swi6 is a significant contributor to the viscoelastic behavior of the nucleus. We propose that the ensemble mechanics of the nucleus arise from a composite system (individual, HC condensates enmeshed within the chromatin polymer) that has emergent properties, much like what is seen in aqueous foams (e.g., shaving cream)^63^. We suggest that it is this emergent property that provides a mechanism by which these condensates can support nuclear resistance to potentially damaging compression or tension forces. Moreover, this property likely influences the response to forces that can activate changes in the relative amount of HC, specifically of the H3K9me2/3-associated constitutive HC predominantly found at the nuclear periphery^64,65,20^ that may have large-scale consequences on gene silencing, prevention of cellular differentiation^66^, and genome-protective nuclear softening. We also suggest that this composite system likely applies to other biomolecular condensates, particularly those that are enmeshed in biological polymers such as the cytoskeleton.

As described previously for HP1-α and Swi6^22,23,67^, our observations support a model in which Swi6 exhibits liquid-like properties. As telomere clustering is compromised upon loss of Swi6, expression of the *swi6-sm1* allele, or treatment with 1,6-hexanediol, we suggest that the formation of Swi6-dependent condensates drives coalescence of telomeric HC, as has been suggested recently for telomeres in other contexts^45,46,49^. The phenotypic mimicry of the *swi6-sm1* allele supports previous work showing that the N-terminus of HP1-α/Swi6 serves an essential role in the formation of phase-separated domains^22,55^. As the *swi6-sm1* allele also confers a silencing defect despite normal recruitment to H3K9me regions^53^, it also offers a potential link between HP1-α/Swi6 condensation and transcriptional silencing, which will require further study.

While recent work affirms that degradation of HP1-α also leads to softer nuclei in mammalian cells suggested to arise from crosslinking of H3K9me2/3 nucleosomes^21^, our findings specifically suggest an alternate model for the underlying mechanism, namely that the ability of the mobile phase of HP1-α/Swi6 to undergo condensation is essential to its contribution to nuclear stiffness. Previously, we showed that the tethering of chromatin to the nuclear membrane acts as a mechanical scaffold for the nucleus, and that the nucleus behaves as a viscoelastic material, exhibiting both liquid and solid characteristics that are time-dependent^12^. This dual nature allows the nucleus to resist deformation (elasticity/stiffness) while allowing plasticity on longer time scales to enable chromatin flow within the nucleus (viscosity). The increased nuclear viscosity associated with expanding H3K9 HC domains in cells lacking Epe1 in our *in vitro* measurements further implies reduced chromatin flow under these conditions. In this context, expansion of small HC domains within the chromatin polymer mesh may promote a more elastic, solid behavior on the minutes timescale (Figure 5D). *In vivo*, however, loss of Epe1 has little effect on local nuclear deformations due to microtubule shearing force while further expansion of HC with concomitant loss of Mst2 leads instead to increased nuclear deformability, suggesting that the loss (and not the gain) of the mobile, condensed Swi6 pool dominates the local nuclear behavior in response to intrinsic cytoskeletal forces. This mechanical transformation has potential implications for physiologic states where cells exhibit gains in repressive H3K9 methylation marks, for example in response to critical stretching^20,65^. Here, our data suggest that increases in H3K9 methylation not only act to repress transcription but could also increase nuclear stiffness and decrease nuclear viscosity when at moderate levels. However, if pronounced, high levels of H3K9 methylation could instead cause nuclear softening should it outpace the available pool of unbound HP1-α/Swi6 in the HC-associated condensate – suggesting a “Goldilocks” balance required to retain both bound and mobile pools of HP1-α/Swi6 to impart optimal nuclear stiffness.

Loss of HP1-α/Swi6 in the condensate, either through complete deletion of Swi6 or by expression of the *swi6-sm1* allele, destabilizes the nucleus, reducing both its ability to resist deformation and impede chromatin flow. This effect is similar to the softer, less viscous nuclei that lack the H3K9 methyltransferase, Clr4, which controls recruitment of HC-condensing proteins (including Swi6). The similarity in the mechanical consequences of disrupting Clr4 or Swi6 is consistent with a model in which the specific properties of Swi6 associated with HC are necessary for the mechanical effects of H3K9me2/3 HC to manifest rather than arising through a direct effect of H3K9 methylation levels. This finding also excludes a quantitative contribution of other H3K9me2/3-binding HC proteins such as Chp1 and Chp2 to nuclear mechanics, singling out Swi6 as both necessary and unique.

It is important to note that the biological importance of the ability of HP1-α to undergo condensation *in vitro* on its *in vivo* function has been questioned, for example the failure of HP1-α over-expression to increase the size of heterochromatic domains as predicted for simple liquid-liquid phase-separation systems^68^. However, the complexity of feedback within HC is rife with paradox, for example the requirement for siRNA transcription to drive chromatin compaction and silencing in fission yeast^69^. Indeed, Swi6 recruits the H3K9me2/3 demethylase Epe1 almost exclusively into HC domains^70^, which would be expected to decrease available H3K9me2/3 marks and thereby delocalize H3K9me2/3-engaged Swi6 molecules, likely short-circuiting the ability of elevated Swi6 levels to drive HC spreading. Moreover, the binding of HP1-α/Swi6 to H3K9me2/3 and its subsequent compaction of HC involves an integrated set of activities driven by its (1) chromodomain avidity, (2) dimerization, (3) engagement of nucleosome arrays, and (4) its condensation properties, all of which make it challenging to arrive at mechanistic conclusions without clear separation-of-function mutants or through engineered systems^68^. We believe this study, and the use of the *swi6-sm1* allele, provides an additional context in which condensation of HP1-α/Swi6 has a critical function – namely in supporting nuclear mechanics.

Taken together, our results suggest that the mobile, condensed phase of HP1-α/Swi6 at HC domains contributes stiffness to the nucleus, similar to the ability of phase-separated droplets to provide structure to a polymer mesh in which they are embedded^71^. Thus, we have extended the roles for biomolecular condensation *in vivo* beyond cellular compartmentalization to include mechanical functions at the level of micron scale organelles, expanding our perspective on the multi-faceted functionality of condensates in cellular biology.

## ACKNOWLEDGEMENTS

We thank members of the King, Mochrie, and Lusk laboratories for helpful discussions, commentary, and troubleshooting, in particular, Hao Yan, Elisa Rodriguez, Andrea Nguyen, and Patrick Lusk. We thank Saikat Biswas for his initial support in creating the PAmCherry-*swi6-sm1* strain for single molecule measurements. We thank the Yeast Genome Resource Center at Osaka University and the many investigators who have deposited strains at this resource. This work was supported by the NIH Pre-doctoral Program for Cellular and Molecular Biology (T32GM00722343), the NSF (CMMI-1334406), the Physical and Engineering Biology Program at Yale, a National Science Foundation Understanding Rules of Life Award (1921677) to JSB and KR, and an NIH award R35GM137832 to KR.

## AUTHOR CONTRIBUTIONS

Conceptualization, J.W. and M.K.; Data curation, J.W., I.S., G.R., and H.N.; Formal analysis, J.W., I.S., H.N., Z.C., and M.K.; Funding acquisition, J.B., K.R., S.M. and M.K.; Investigation, J.W., I.S., S.S., Z.C., G.R., Y.H., and M.K.; Methodology, J.W., I.S., Z.C., J.B, K.R., S.M., and M.K.; Project administration, M.K.; Resources, J.B., K.R., S.M. and M.K.; Software, J.W., I.S., Z.C., H.N., and J.B.; Supervision, J.B., K.R., S.M. and M.K.; Validation, J.W., I.S., and M.K.; Visualization, J.W., I.S., G.R., K.R., and M.K.; Writing – original draft, J.W. and M.K.; Writing – review & editing, J.W., I.S., S.S., Z.C., G.R., H.N., Y.H., J.B., K.R., S.M., and M.K.

## STAR METHODS

## RESOURCE AVAILABILITY

### Lead contact

Further information and requests for resources and reagents should be directed to and will be fulfilled by the Lead Contact, Megan King.

### Materials availability

All unique/stable reagents and strains generated in this study are available from the Lead Contact without restriction.

### Data and code availability

The code generated during this study (3DMembraneReconstruction, Swi6/*swi6-sm1* diploid heterochromatin domain detection and analysis, TweezersAnalysis, and NuclearFluctuations) is available on GitHub at https://github.com/LusKingLab/.

## METHOD DETAILS

### Cell culture and strain generation

The strains used in this study are listed in the Key Resources Table (see also reference^72^). *S. pombe* cells were grown and maintained in standard cell culture conditions at 30°C^73^. Genetic knock-outs were made by gene replacement of the open reading frames with the kanMX6^74^, hphMX6, or natMX6 cassettes^75^. C- and N-terminal GFP tagging was performed with pFa6a-GFP-kanMX6^74^ or pFa6a-natMX6-Pnmt41^74^ respectively; the pFa6a-mCherry-kanMX6 cassette was also used as a template for C-terminal mCherry tagging^76^. To edit PAmCherry-Swi6 to generate PAmCherry-Swi6-sm1, we used a CRISPR-Cas9 based approach. sgRNAs targeting the N-terminus of Swi6 were generated and transformed as previously described in Torres-Garcia *et al*.^77^ along with a gene block (Twist Biosciences, CA) that contains the sm1 allele mutations (D21G, ΔE74) and synonomous substitutions that destabilize the sgRNA binding site. The resulting *S.pombe* colonies were screened using Sanger sequencing to confirm the desired mutation^77^. All strains generated by cassette integration were confirmed by PCR and/or DNA sequencing. Strains made through genetic crosses were confirmed by the segregation of selection markers and/or by the presence of the designated fluorescently-tagged protein.

### Standard live-cell microscopy

*S. pombe* strains were grown in YE5S plus 250 mg/L adenine to log phase (OD_600_ 0.5-0.8). Cells were mounted on agarose pads (1.4% agarose in EMM5S) and sealed with VALAP (1:1:1, vaseline:lanolin:paraffin). Cells treated with MBC were incubated on agar pads with 50 mg/mL MBC for 10 min before imaging. Cells treated with 1,6-hexanediol were incubated with final concentrations of 5% 1,6-hexanediol for 15 or 30 min as indicated in the legend and imaged directly thereafter. Live-cell images were acquired on a Deltavision Widefield Deconvolution Microscope (Applied Precision/GE Healthcare) with a CoolSnap HQ2 CCD camera (Photometrics) or an Evolve 512 EMCCD camera (Photometrics). For Swi6-GFP time-lapse imaging, single mid-plane z-slices with 0.15-sec exposure time were taken every 5 sec with the CCD camera. For telomere/Taz1-GFP imaging, 15 z-slices with 50-ms exposure time and 320-nm spacing were taken with the EMCCD camera. For Swi6/*swi6-sm1*-GFP imaging, 16 z-slices with 10-ms exposure time and 400-nm spacing with taken with the EMCCD camera. For Swi6/*swi6-sm1* diploid imaging, 21 z-slices with 500-ms exposure time and 200-nm spacing with taken with the EMCCD camera. For nuclear contour imaging of Cut11-GFP, 10 z-slices with 20-ms exposure time and 400-nm spacing were taken every 2.5 s for 5 min with the CCD camera. For GFP-Swi6 imaging, 20 z-slices with 0.1-sec exposure time and 200-nm spacing with taken with the EMCCD camera. For Chp1-GFP imaging, 20 z-slices with 0.2-sec exposure time and 200-nm spacing were taken with the CCD camera.

### Heterochromatin shape and telomere analysis

To characterize the shape dynamics of Swi6-GFP foci, single mid-plane z-slices were read in MathWorks MATLAB R2016b (Natick, MA, USA), binarized with a pixel threshold value of 0.8, and focus boundary x and y values were found using the function bwboundaries. X and y values for each Swi6-GFP focus shape was plotted using the MATLAB fill function.

For Taz1-GFP and cell length analysis only live, interphase cells that had completed cytokinesis were considered. Maximum intensity projections were displayed as described in the Figure legends. Images were analyzed in ImageJ (Fiji;^78^). To calculate the number of Taz1-GFP foci per cell, maximum Z-projections were generated and total number of Taz1-GFP foci per cell were counted. For statistical comparison, percentage of cells treated with 1,6-hexanediol, *swi6Δ* cells, and *swi6-sm1* cells were individually compared to wild-type cells for each cell length (a proxy for cell cycle stage) using Fisher’s exact test.

### Three-dimensional heterochromatin foci analysis

GFP-Swi6, Swi6-GFP, and *swi6-sm1*-GFP cross-sectional foci areas and intensities were measured using three-dimensional reconstructions of GFP foci with a custom MATLAB software as previously described^56^. Foci were grouped by cell using a minimum separation distance of 40-50 pixels from the center of the grouping; this adequately grouped together foci of the same nucleus while excluding foci from nearby, neighboring cells. Z-stack images for each focus were individually fit with a three-dimensional Gaussian function to determine maximum focus width in both x and y-dimensions as well as foci integrated intensity. Foci intensity vs. cross-sectional area plots and histograms were generated using the MATLAB functions dscatter^79^ and scatterhist. For statistical comparison, Swi6-GFP (wild type) and *swi6-sm1-*GFP foci areas and intensities were compared using the Kolmogorov-Smirnov test.

### Single-molecule tracking photoactivation localization microscopy (spt-PALM)

Yeast strains containing a copy of *swi6-sm1*-PAmCherry under the control of the native Swi6 promoter were grown in standard complete YES media (US Biological, cat. Y2060) containing the full complement of yeast amino acids and incubated overnight at 32 °C. The seed culture was diluted and incubated at 25 °C with shaking to reach an OD_600_ ∼0.5. To maintain cells in an exponential phase and eliminate extranuclear vacuole formation, the culture was maintained at OD_600_ ∼0.5 for 2 days with dilutions performed at ∼12-hour time intervals. To prepare agarose pads for imaging, cells were pipetted onto a pad of 2% agarose prepared in YES media, with 0.1 mM N-propyl gallate (Sigma, cat. P-3130) and 1% gelatin (Millipore, cat. 04055) as additives to reduce phototoxicity during imaging. *S. pombe* cells were imaged at room temperature with a 100× 1.40 NA oil-immersion objective in an Olympus IX-71 inverted microscope. First, the fluorescent background was decreased by exposure to 488-nm light (Coherent Sapphire, 200 W/cm^2^ for 20 – 40 s). A 406-nm laser (Coherent Cube 405-100; 102 W/cm^2^) was used for photo-activation (200-ms activation time) and a 561-nm laser (Coherent-Sapphire 561-50; 163 W/cm^2^) was used for imaging. Images were acquired at 40-ms exposure time per frame. The fluorescence emission was filtered with Semrock LL02-561-12.5 filter and Chroma ZT488/561rpc 488/561 dichroic to eliminate the 561-nm excitation source and imaged using a 512 × 512-pixel Photometrics Evolve EMCCD camera.

### Single-molecule trajectory analysis with SMAUG algorithm

Recorded *swi6-sm1*-PAmCherry single-molecule positions were detected and localized with 2D Gaussian fitting with home-built MATLAB software as previously described and connected into trajectories using the Hungarian algorithm^80,81^. These single-molecule trajectory datasets were analyzed by a non-parametric Bayesian framework to reveal heterogeneous dynamics^58^. This SMAUG algorithm uses non-parametric Bayesian statistics and Gibbs sampling to identify the number of distinct mobility states, *n*, in the single-molecule tracking dataset in an iterative manner. It also infers the weight fraction, *π*_i_, and effective diffusion coefficient, *D*_i_, for each mobility state (*i* = 1,…,*n*), assuming a Brownian motion model. We ran the algorithm over >10,000 iterations to achieve a thoroughly mixed state space. The state number and associated parameters were updated in each iteration of the SMAUG algorithm and saved after convergence. The final estimation shows the data after convergence for iterations with the most frequent state number. Each mobility state, *i*, is assigned a distinct color, and for each saved iteration, the value of *D_i_* is plotted against the value of *π_i_*. The distributions of estimates over the iterations give the uncertainty in the determination of *D_i_*. Furthermore, the transition probabilities (e.g., Figure 3D,E) give the average probability of transitioning between states from one step to another in any given trajectory.

### Swi6/*swi6-sm1* diploid heterochromatin domain detection and analysis

Swi6 HC domains were detected in 3D images in both GFP and mCherry channels independently using custom-written MATLAB code that finds irregular fluorescent blobs regardless of their shape (https://github.com/LusKingLab/Spot_Detection_n_Colocalization). First, the code identifies all pixels that are brightest within sub-volumes of a given size. Next, it calculates ratio of the mean intensities of “central” pixels (*i.e.*, pixels within a given radius *R_cent_*, from the brightest pixel) versus “edge” pixels (*i.e.*, pixels within a spherical shell of a given thickness *d_shell_* ith *R_edge_* sphere radius). If the ratio is above a given threshold, the code keeps the identified spot and calculates its 3D position as a intensity-weighted centroid of pixels within *R_fit_* radius, and its peak intensity as a mean intensity of pixels within *R_fit_* radius. After spot identification, individual GFP and mCherry spots were paired if the peak positions were within 3 pixel distance. These paired spots were used to compare intensities between GFP and mCherry signals and to calibrate channels relative to each other (using data from cells bearing only wild-type Swi6 - Swi6-GFP/Swi6-mCherry cells), and to compare recruitment of Swi6-mCherry versus *swi6-sm1*-mCherry.

To analyze data for centromeric HC domains versus other HC domains, domains detected in the GFP channel were grouped into nuclei based on spatial proximity. Next, for each nucleus, the nuclear background intensity was determined as a median intensity value for all pixels within *R_nuc_* radius. Then, for each nucleus with at least two spots (*n_spots_* > 2), the peak intensity of each spot was assigned to the “bright” (*i.e.*, centromeric), “intermediate”, or “dim” class. Note that in this “by nucleus” definition a bright spot from one nucleus might be dimmer than an intermediate spot from another. The peak intensities of the classified spots in the GFP channel were then compared to their paired intensities in the mCherry channel. To determine the calibration coefficient between GFP and mCherry channels, data from “centromeric” (“bright”) domains below an intensity threshold (to avoid the influence of few very bright domains) after correction for different exposure time were fitted by *ImCherry* = a\**<u>I_GFP_</u> + b*, resulting in calibration coefficient *a* = 0.4412, that was then used in the subsequent analysis.

### Nuclear isolation

Nuclei were isolated as previously described in ^12^. Briefly, strains were grown to log phase in rich media (YE5S) at 30°C and diluted into 1 liter YE5S for overnight growth. Cells were harvested the following day at an OD_600_ ∼0.8 and incubated in 100 mM Tris pH 9.4 and 10 mM DTT for 10 min at 33°C to prepare for spheroplasting. Cells were spheroplasted in 0.4 mg/mL zymolase (MP Biomedicals), 0.6 mg/mL lysing enzymes (Sigma), 350 µL beta-glucuronidase (MP Biomedicals) and 5 mM DTT in 1.1 M sorbitol at 33°C for 2-3 hours. Spheroplasts (nuclei) were isolated from cells by centrifugation using a series of three density-step sucrose gradients, and aliquots were flash frozen in liquid N_2_ and stored at -80°C. To thaw for use, one aliquot was dialyzed overnight in 500 mL dialysis buffer (80 mM PIPES, 5% DMSO, 2 mM MgCl_2_, 1 mM EGTA, 500 mM sucrose).

### Force spectroscopy oscillation assay

Force spectroscopy was performed on isolated nuclei as described previously in Schreiner *et al.*^12^. In a flow cell, isolated *S. pombe* nuclei were trapped and fixed via electrostatic interactions to poly-ornithine-coated silica beads by micromanipulation using optical tweezers. Optical trapping and interferometry position detection was achieved using a single-beam gradient focus from a 3-watt 1064 nm ND:YAG laser. The focused laser was used to trap beads and nuclei within the flow cell, which was mounted onto a piezo-controlled stage with nanometer precision. To visualize each nucleus, an epifluorescence microscope was coupled with the optical tweezers, equipped with a 470 nm excitation source (blue light emitting diode). Each nucleus (using the optical tweezers) was affixed on its right side to a 5.2 μm bead adherent to a glass coverslip, which elevates the nucleus off the coverslip and provides a stationary structure upon which forces can be applied. Next, a smaller 1.2 μm bead (using the optical tweezers) is affixed to the left (free) side of the nucleus, creating a bead-nucleus-bead sandwich. While holding the smaller bead within the optical trap, the large bead and coverslip—attached to a piezostage—is respectively translated away and towards the optical trap by a pre-determined displacement (Δ*x*_piezostage_) to exert respective tension or compression on the sandwiched nucleus. The resulting displacement of the small bead in the optical trap (Δ*x*_bead_) is measured by its deflection within the trap using back focal plane interferometry via an infrared-enhanced quadrant photodiode. The difference in the two displacements is equal to the displacement of the nucleus (Δ*x*_nucleus_). To exert compression and tension forces, the stage is driven sinusoidally with an amplitude range of 60-100 nm (physiologic length scales), at discrete frequencies of 0.01-2 Hz.

### Force spectroscopy data analysis

To determine the effective stiffness of the nucleus (*k_nucleus_*), the system was treated as a spring that adheres to Hooke’s law (*F = k_nucleus_*Δ*_nucleus_*), where Δ*_nucleus_* is the displacement of the nucleus (either in extension or compression) and *F* is the force imparted to the nucleus from the oscillating stage. Δ*_nucleus_* was calculated as the difference between the piezo stage displacement (Δ*_piezostage_*) and the displacement of the small bead from the center of the trap (Δ*_bead_*). The force *F* on the nucleus is equal and opposite to the force on the bead in the trap, therefore determining the stiffness of the trap allowed us to calculate the force acting upon the bead and thus the force acting upon the nucleus. The inherent stiffness of the optical trap was determined for each experiment by fitting the measured power spectral density of a trapped 1.2 µm bead with a Lorentzian function to extract the corner frequency^82^. The trap stiffness is the ratio of this corner frequency to the bead’s friction coefficient. We subjected each nucleus to multiple rounds of compression and extension at different frequencies (0.01 – 2 Hz), enabling us to observe any time-dependent mechanical properties. We then fit the effective stiffness data with a Maxwell viscoelastic model to extract the elastic (spring) and viscous (dashpot/dampener) components, *K* and η respectively, using the corresponding fitting function for the effective stiffness:

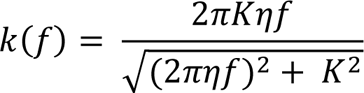

### Nuclear envelope deformation analysis

NE contour fitting and deformation time analysis were performed as previously described in ^12^ and ^56^. Briefly, our contour fitting algorithm reconstructed the three-dimensional nuclear shape with sub-pixel resolution from a z-stack of 10 two-dimensional images by optimizing NE shape to simultaneously maximize the total intensity of the nuclear envelope across the z-stack and minimize the NE’s total curvature. To avoid the effects of aberration and difficulties with image noise at the nuclear caps in the z-dimension, we limited our deformation analysis to a band 45 degrees above and below the center/“equator” of the nucleus (Figure 6D and Figure S7a). After the 3D shape reconstruction at each time point (denoted by the radial distance from the nuclear center r(θ, ϕ, *t*) at vertex angle (θ, ϕ) at frame *t*), a time-average shape *R_av_*(*θ*, *φ*) = <*r*(*θ*, *φ*, *t*)>*_t_* was calculated and shape deformations were detected by deviation of the radial distance from a time-averaged radial distance at that same (*θ*, *φ*) position. The average, or root mean squared, deformation size was calculated according to:

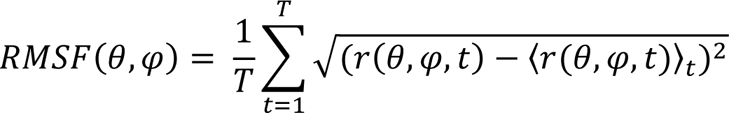

Where *T* is the total number of frames, r(θ, ϕ*, t*) is the radial distance from the nuclear center at vertex angle (θ, ϕ) at frame *t* and < r(θ, ϕ*, t*) >*_t_* denotes the time-averaged radial distance. Individual deformations were tracked over time using 2D particle tracking, assigning longer deformations (> 10 frames/25 s) to those induced by microtubules and shorter deformations arising from baseline thermal activity. Peak height, formation (rise) time, and resolution (decay) time were determined by fitting each deformation trajectory to a single, asymmetric triangle waveform.

## QUANTIFICATION AND STATISTICAL ANALYSIS

Statistical details (statistical tests used, exact value for n, what n represents, definition of center, and dispersion and precision of measures) for each experiment can be found in the Figure legends. Microsoft Excel (Redmond, WA, USA) was used to tabulate values, GraphPad Prism 9 (La Jolla, CA, USA) was used to create simple plots (Figure 1D, Figure 2E, Figure 5C, D, E, F, Figure S1C, D, Figure S3F, Figure S4, and Figure S6) and perform statistical analysis, and MathWorks MATLAB R2016b (Natick, MA, USA) was used for all other analysis and plotting. * p < 0.05; ** p < 0.01; *** p < 0.001; **** p < 0.0001 by Fisher’s exact test, ordinary one-way ANOVA with Dunnett’s multiple comparisons test, unpaired t-test, Kolmogorov-Smirnov test, or Mann-Whitney U test.

## SUPPLEMENTAL ITEM TITLES

**Table S1. Diffusion coefficients and weights from Figures 3C and 7D**

