## Supplemental Figures S1-S8 and Table S1 for "The condensation of HP1-α/Swi6 imparts nuclear stiffness"

**Williams et al.; SUPPLEMENTAL FIGURES and Table S1**

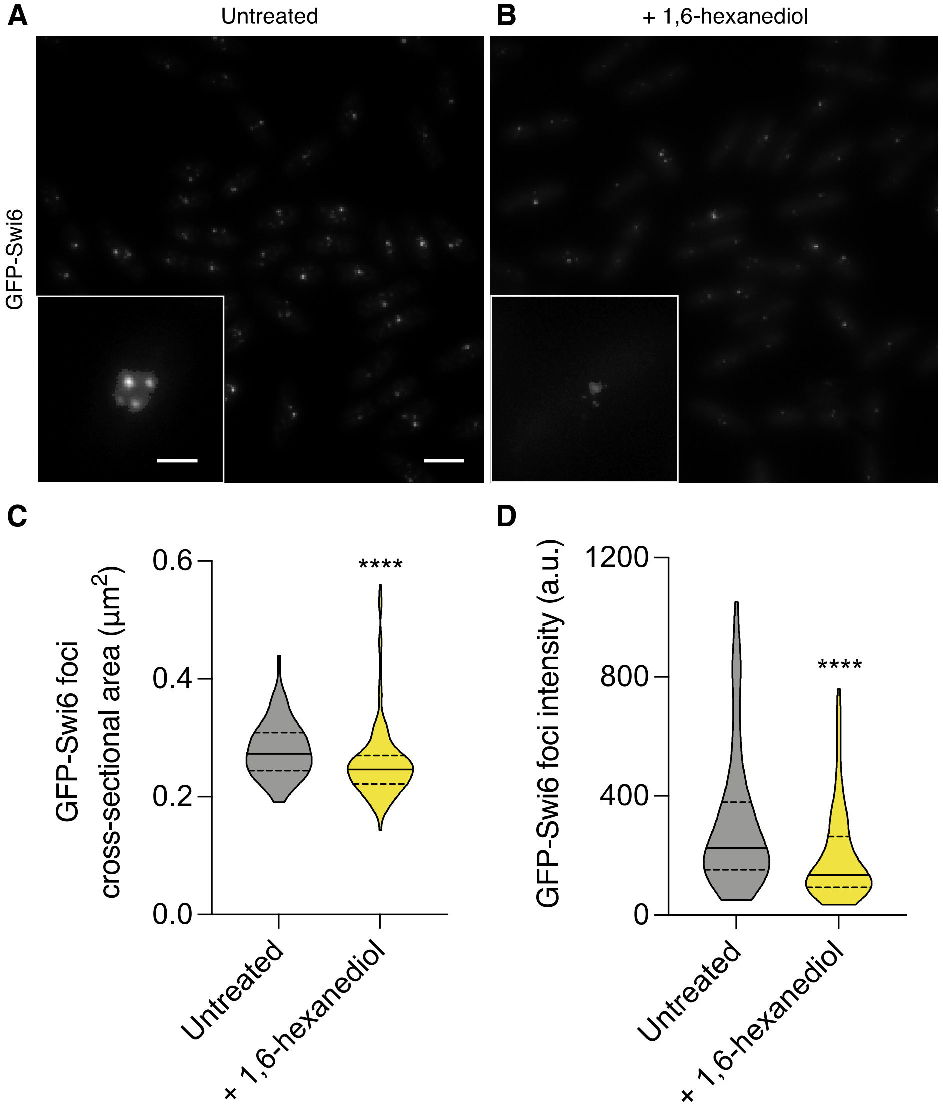


**Figure S1. GFP-Swi6 foci decrease in size and intensity when cells are treated with 1,6-hexanediol**

(A) N-terminally-tagged GFP-Swi6 labels HC foci in live cells. Fluorescence maximum intensity projection of wild-type *S. pombe* cells expressing GFP-Swi6. Scale bars, 6 µm and 2 µm (insert).

(B) Treatment with 5% 1,6-hexanediol for 15 min results in diminished GFP-Swi6 foci signal.

(C,D) Cells treated with 1,6-hexanediol (yellow, n = 57 cells; 145 foci) contain smaller and dimmer GFP-Swi6 foci as compared to untreated cells (gray, n = 48 cells; 169 foci). Area and intensity measurements were obtained by 3D foci reconstruction (see Methods and reference[56]) **** $p < 0.0001$ by Kolmogorov-Smirnov test.

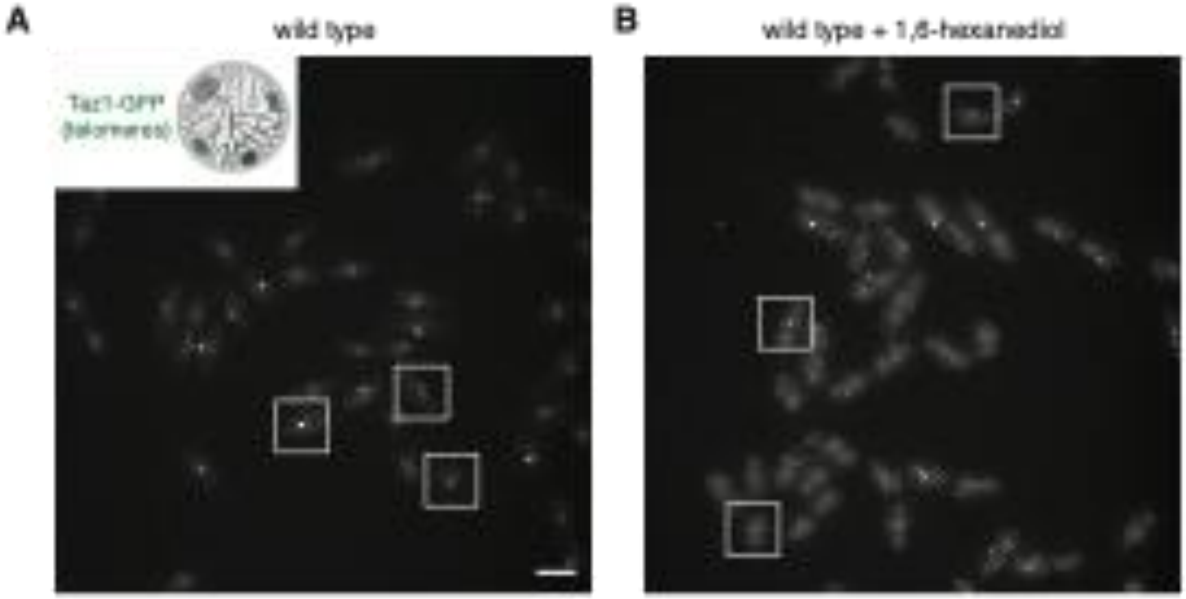


**Figure S2. Labeling of telomeres with Taz1-GFP and the de-clustering effect of treatment with 1,6-hexanediol**
(A) Under normal conditions, the six fission yeast telomeres typically cluster into ~1-3 foci during interphase (boxes with magnifications below). Fluorescence maximum intensity projection of wild-type *S. pombe* cells expressing Taz1-GFP to label telomeres. Scale bar, 6 μm.
(B) Treatment of wild-type cells with 5% 1,6-hexanediol for 30 min leads to telomere de-clustering, as the number of Taz1-GFP foci increases to ~4-6 per nucleus (see quantification in Figure 1C,D).

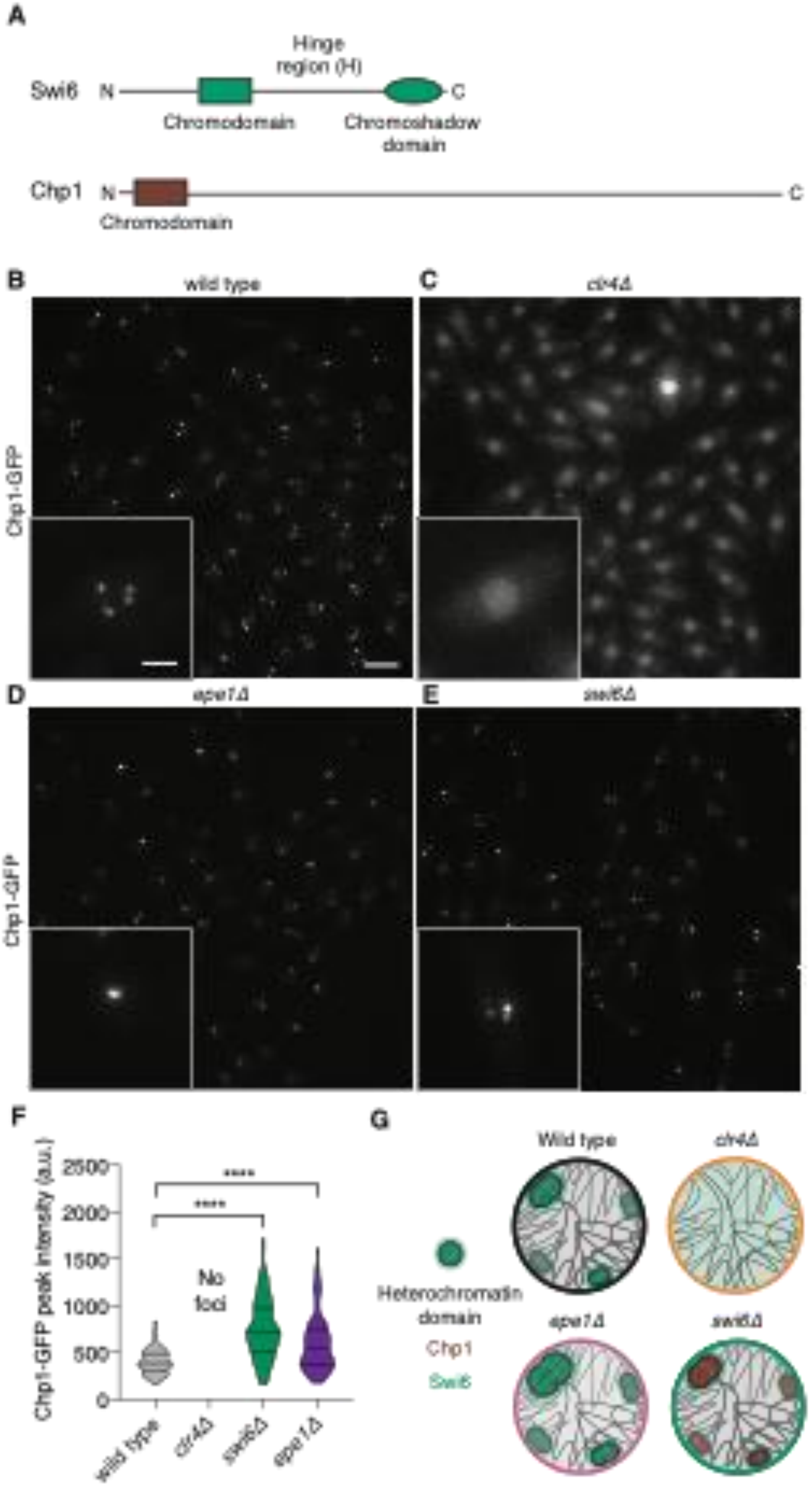

A
Hinge region (H)
Swi6
N
C
Chromodomain
Chromoshadow domain
Chp1
N
C
Chromodomain
B
wild type
C
clr4Δ
Chp1-GFP
D
epe1Δ
E
swi6Δ
Chp1-GFP
F
Chp1-GFP peak intensity (a.u.)
2500
2000
1500
1000
500
0
****
****
No foci
wild type
clr4Δ
swi6Δ
epe1Δ
G
Wild type
clr4Δ
Heterochromatin domain
Chp1
Swi6
epe1Δ
swi6Δ

**Figure S3. Swi6 competes for H3K9me3 binding sites with other chromodomain proteins and plays a unique role in domain clustering**

(A) Protein diagrams of Swi6 and Chp1, HC proteins in fission yeast (drawn to scale). Swi6 (328 aa) consists of an unstructured N-terminal domain, a chromodomain, an unstructured hinge domain, and a chromo-shadow domain. In contrast, Chp1 (960 aa) contains only a chromodomain with only a much small N-terminal domain and extended C-terminal domain. Fluorescence maximum intensity projections of Chp1-GFP in (B) wild type, (C) Suv39/Clr4-deficient, (D) Epe1-deficient, and (E) Swi6-deficient *S. pombe* cells. Insets show HC foci from one nucleus. Scale bars, 6 μm and 2 μm.

(F) Chp1-GFP intensity at HC domains increases in response to Swi6 or Epe1 deletion while no foci are apparent with loss of H3K9 methylation upon deletion of Clr4. Quantification of Chp1-GFP peak intensity for wild type (n = 117 foci), *epe1Δ* (n = 84 foci), and *swi6Δ* (n = 91 foci) cells. **** $p < 0.0001$ by ordinary one-way ANOVA with Dunnett's correction for multiple comparisons.

(G) Schematic representation of H3K9me-associated HC domains in each genotype. Wild-type cells contain both Chp1 and Swi6 (and Chp2), and *clr4Δ* cells have delocalized Chp1 and Swi6. Without H3K9me demethylation (*epe1Δ*), HC domains spread, allowing for more HC to be bound by Chp1 (and Swi6). When Swi6 is removed, more Chp1 is able to bind existing HC foci, resulting in increased Chp1-GFP foci intensity.

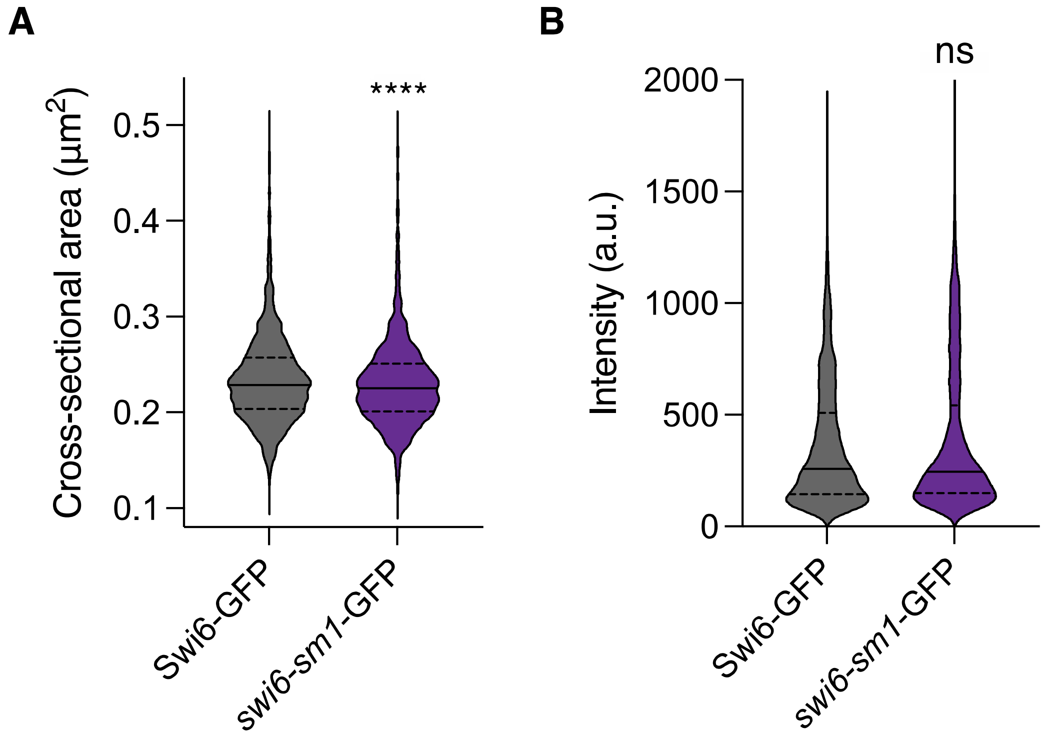


**Figure S4. Aggregate population distributions of *swi6-sm1*-GFP and Swi6-GFP foci are similar in range of size and intensity by 3D foci reconstructions**

(A) Cross-sectional area and (B) intensity of Swi6-GFP (n = 2500 foci) and *swi6-sm1*-GFP (n = 2402 foci) labeled HC domains. **** $p < 0.0001$ by Mann-Whitney U test.

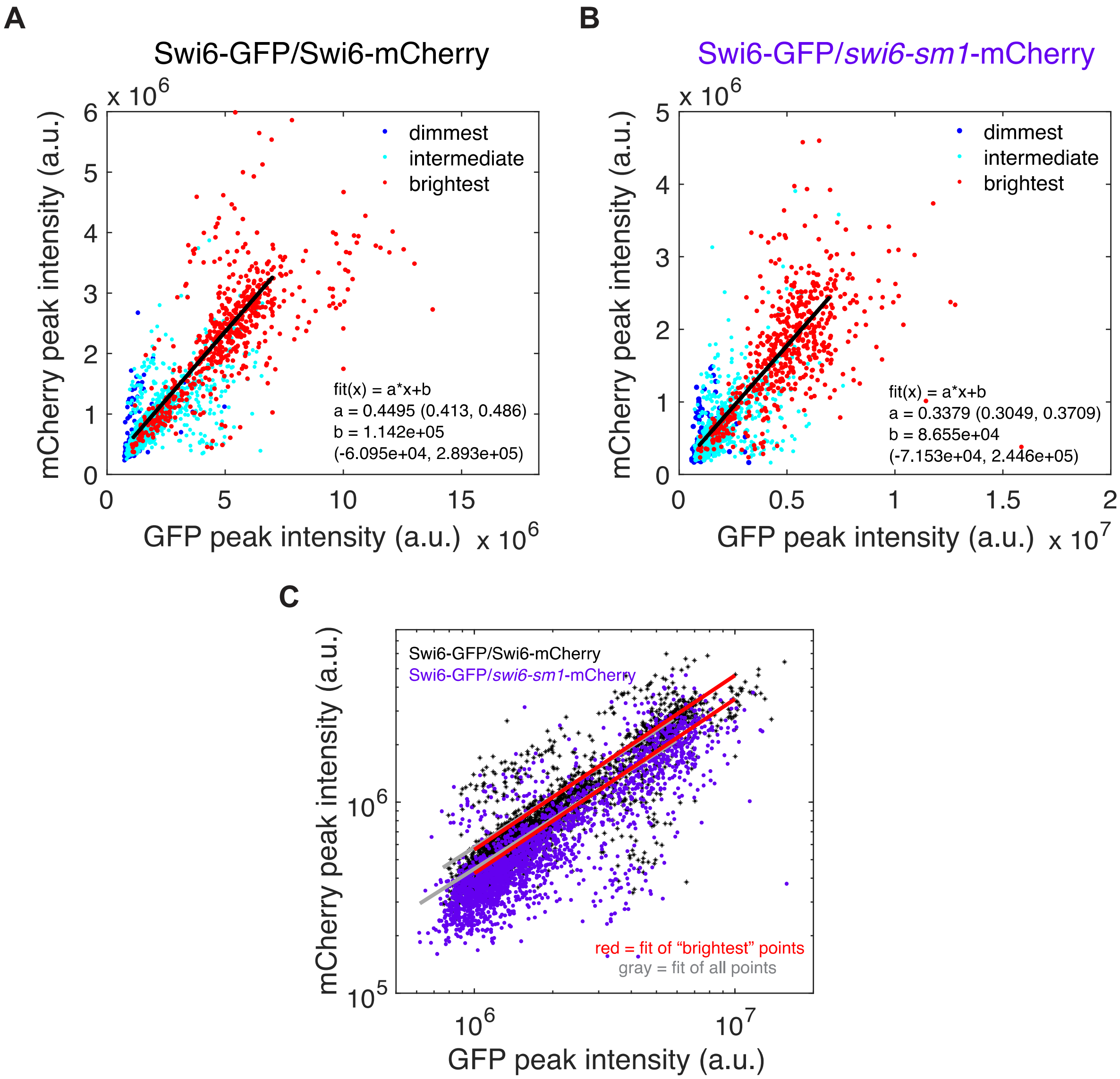


**Figure S5. *swi6-sm1*-mCherry is depleted relative to WT Swi6-mCherry at HC foci in diploid cells also expressing WT Swi6-GFP**

(A) Scatter plot of Swi6 foci intensities in diploid cells from nuclei with ≥ 3 foci, shown as mCherry signal versus GFP signal for Swi6-GFP/Swi6-mCherry cells (n = 584 nuclei; 2,648 foci) and (B) Swi6-GFP/*swi6-sm1*-mCherry cells (n = 538 nuclei; 2,327 foci). Swi6 foci are separated by "brightest" (red), "intermediate" (cyan), and "dimmest" (blue) intensity per nucleus. Black lines are best fit using "brightest" foci of a linear model y(x) = a*x+b, with a = 0.4495, b = 1.142 x $10^5$ (Swi6-GFP/Swi6-mCherry) and a = 0.3379, b = 8.655 x $10^4$ (Swi6-GFP/*swi6-sm1*-mCherry), respectively.

(C) mCherry versus GFP log-log scatter plot of Swi6 foci intensities with linear fit based on "brightest" foci only (red); also shown is the linear fit based on all foci (gray): a = 0.4412 (95% confidence bounds: 0.4296, 0.4528), b = 3.302 x $10^4$ (1,534, 6,451 x $10^4$) (Swi6-GFP/Swi6-mCherry) and a = 0.3313 (0.3224, 0.3403), b = 2.671 x $10^4$ (2,501, 5.092 x $10^4$) (Swi6-GFP/*swi6-sm1*-mCherry).

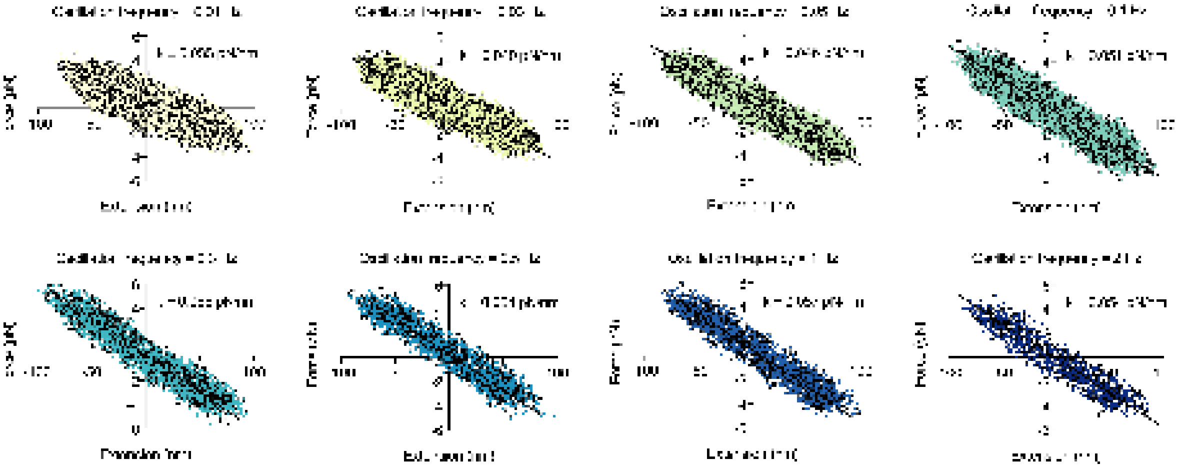

**Figure S6. Examples of force versus extension curves for a single wild-type nucleus**

A single nucleus undergoes extension and compression in a series of consecutive oscillation frequencies ranging from 0.01 – 2 Hz. The resulting force experienced by the nucleus is plotted as a function of nuclear extension. The data display linear relationships (black lines) over each individual oscillation; the slope (k) is equivalent to the spring constant, or stiffness, of the individual nucleus at that oscillation frequency. Over the series of oscillations, the same nucleus displays reduced stiffnesses at the slowest oscillation speeds, typical of a viscoelastic material.

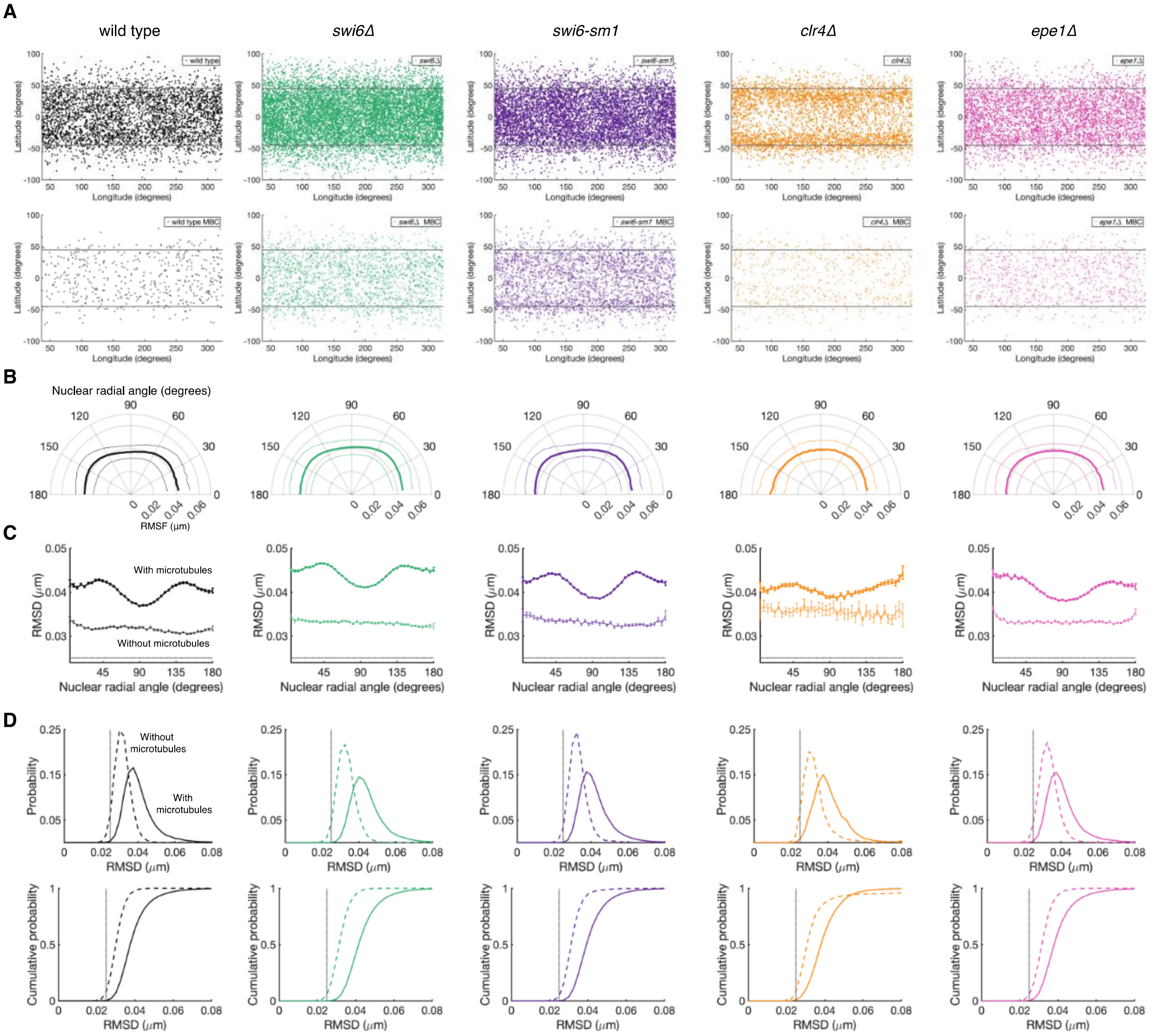


**Figure S7. *In vivo* nuclear deformation analysis by genotype**

(A) For each genotype population, individual nuclear deformations are diffusely detected across the nuclear surface (top row, solid points), except for *clr4Δ* nuclei, which exhibit a depletion of normal deformations at the nuclear "equator." Deformations are plotted with respect to nuclear latitude (y-axis) and nuclear longitude (x-axis). Gray lines are +45 and -45 degrees designate the region included for deformation detection. In the absence of MT dynamics (bottom row, circles), deformations are still detected at diffuse positions across the nuclear surface.

(B) The root mean squared deformation (RMSD) size as a function of nuclear radial angle plotted on a radial axis. Thick line represents the population mean; thin lines represent S.D.

(C) The RMSD size as function of nuclear radial angle plotted on a cartesian/linear axis. For each genotype, mean RMSD size is plotted with and without microtubules present.

Error bars represent S.E.M. Dotted line at 0.025 μm represents the lower limit of deformation detection.

(D) The probability (top row) and cumulative probability (bottom row) of the RMSD size for each genotype with (solid line) and without (dashed line) MT dynamics. Dotted line at 0.025 μm represents the lower limit of deformation detection.

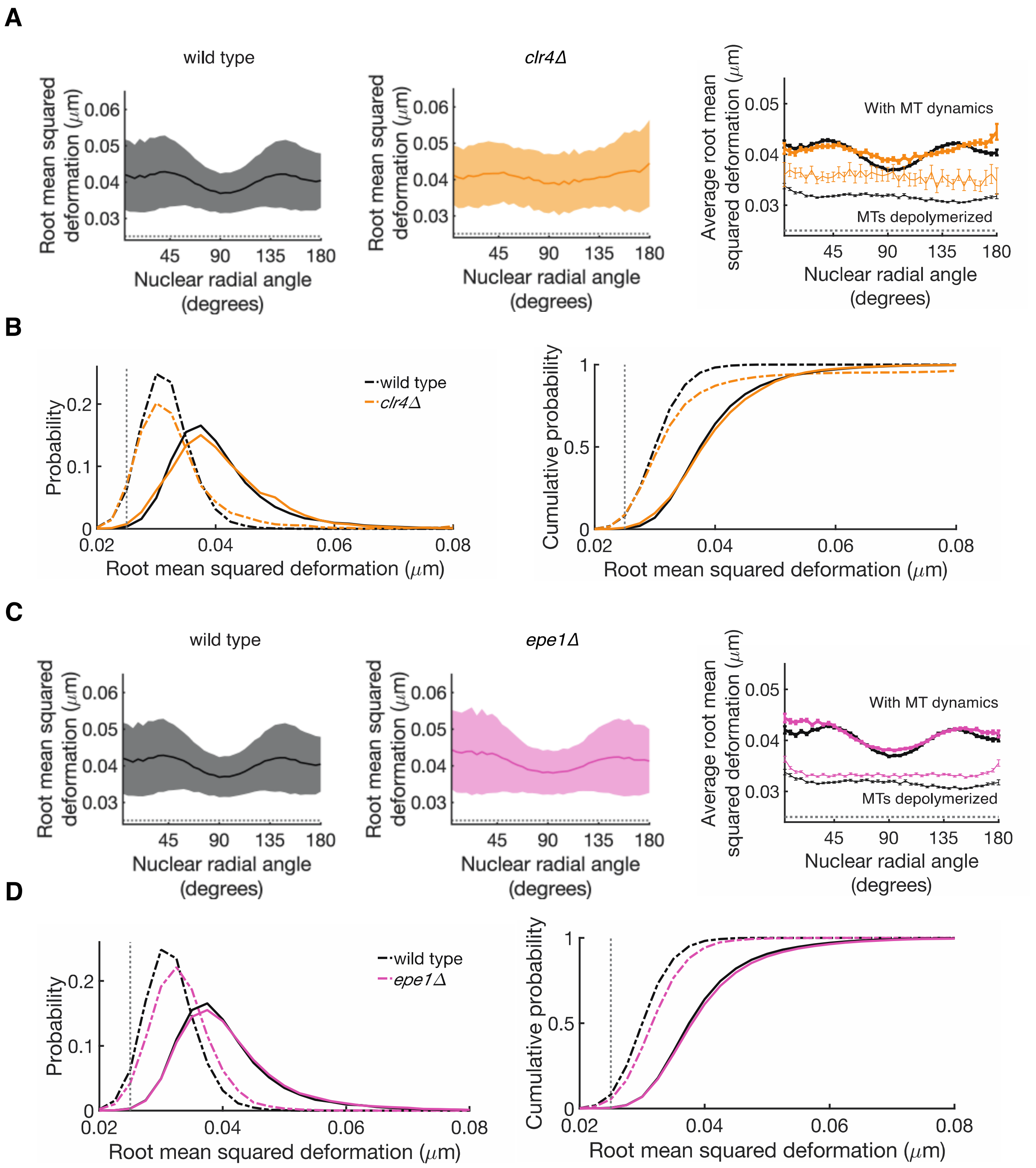


**Figure S8. Loss of the H3K9 methyltransferase Suv39/Clr4 disrupts characteristic MT-induced nuclear deformations, precluding further interpretation; loss of the presumed H3K9me2/3 demethylase KDM2/Epe1 shows no significant nuclear deformation size difference in response to *in vivo* MT forces**

(A) The average, or root mean squared, population deformation size measured with respect to the cross-sectional nuclear radial angle (0 to 180 degrees) displays an aberrant angular profile in *clr4Δ* nuclei, suggesting disruption of the characteristic MT-induced

deformations (population mean, solid line; population standard deviations, band). Wild-type data replotted from Figure 6E.
(B) Without Suv39/Clr4 (orange, n = 66 nuclei; 1,021 deformations), global nuclear deformation size increases. Dashed lines represent recorded nuclear deformations in the absence of MTs (n = 19 nuclei; 261 deformations) as compared to wild-type nuclei (black, replotted from Figure 6E). Dotted line at 0.025 μm represents the lower limit of deformation detection.
(C) The average, or root mean squared, population deformation size measured with respect to the cross-sectional nuclear radial angle (0 to 180 degrees) (population mean, solid line; population standard deviations, band). Wild-type data replotted from Figure 6E.
(D) Without KDM2/Epe1 (pink, n = 215 nuclei; 3,610 deformations), global nuclear deformation size is similar in scale to wild-type nuclei. Dashed lines represent recorded nuclear deformations in the absence of MTs (n = 85 nuclei; 946 deformations) as compared to wild-type nuclei (black, replotted from Figure 6E). Dotted line at 0.025 μm represents the lower limit of deformation detection.

| | *swi6-sm1* | *Swi6 WT* | *2xCD-GST* |
|---|---|---|---|
| Mobility state 1 *(D / μm²/s)* *(Weight)* | 0.010 ± 0.001<br>0.32 ± 0.04 | 0.007 ± 0.001<br>0.23 ± 0.03 | 0.005 ± 0.0003<br>0.16 ± 0.01 |
| Mobility state 2 *(D / μm²/s)* *(Weight)* | 0.02 ± 0.002<br>0.50 ± 0.03 | 0.02 ± 0.003<br>0.32 ± 0.03 | 0.02 ± 0.001<br>0.50 ± 0.01 |
| Mobility state 3 *(D / μm²/s)* *(Weight)* | 0.14 ± 0.02<br>0.15 ± 0.02 | 0.08 ± 0.02<br>0.20 ± 0.02 | 0.06 ± 0.003<br>0.25 ± 0.01 |
| Mobility state 4 *(D / μm²/s)* *(Weight)* | 0.74 ± 0.06<br>0.03 ± 0.01 | 0.51 ± 0.03<br>0.25 ± 0.01 | 0.60 ± 0.02<br>0.09 ± 0.003 |
